# IgD-Expressing Mature B Cells Exhibit Enhanced Sensitivity to Glucocorticoid-Induced Cell Death

**DOI:** 10.1101/2025.10.28.685010

**Authors:** Kais Almohammad, Marc Young, Sabine Vettorazzi, Franziska Greulich, Mahmoud Alkhatib, Jan Tuckermann, Hassan Jumaa, Corinna S. Setz

## Abstract

Glucocorticoids (GCs) regulate diverse physiological processes, comprising metabolism, immune responses, stress adaptation and inflammation. Synthetic GCs are widely used for their powerful anti-inflammatory and immunosuppressive effects, in the treatment of autoimmune diseases, allergies, and inflammation. Here, we investigated the role of the glucocorticoid receptor (GR) in B cell development and survival using both B cell-specific GR-deficient mice and continuous *in vivo* GR agonist treatments. Deletion of the GR in B cells altered splenic B cell subpopulations, increasing follicular and CD21^lo^ B cells and leading to the accumulation of IgM^-^/IgD^-^ B cells. *In vivo* treatment with GR agonists, such as Dexamethasone (Dex) and Prednisolone (Pred), selectively depleted IgD^high^ follicular B cells while enriching IgD^low^ marginal zone B cells. IgD^low^ (IgM^high^) B cells, which were more resistant to glucocorticoid-induced cell death, showed increased expression of IL-10 and genes involved in survival, suggesting a potential regulatory function. *In vitro*, B cell activation via CpG or LPS altered IgM/IgD expression and B cell sensitivity to GR agonists, thereby leading to improved B cell survival and increased plasma cell differentiation. Together, these findings suggest that IgD downregulation and IgM upregulation are critical for B cell survival under GC exposure and that GR agonists promote the enrichment of IgD^low^ (IgM^high^) cells resistant to apoptosis.

## Introduction

Glucocorticoids (GCs) are a class of steroid hormones that play a fundamental role in maintaining homeostasis and regulating numerous physiological processes, including metabolism, immune response, stress adaptation and inflammation (Cruz-Topete and Cidlowski, 2015; De Bosscher and Haegeman, 2009; Eiers et al., 2024; Rhen and Cidlowski, 2005; Sapolsky et al., 2000; Vegiopoulos and Herzig, 2007; Vettorazzi et al., 2022). Produced in the adrenal cortex in response to signals from the hypothalamic-pituitary-adrenal (HPA) axis (Marchi and van Eeden, 2021; Ulrich-Lai and Herman, 2009), GCs are key mediators in adapting to environmental changes and maintaining stability within the body. Their broad regulatory effects encompass a variety of organs and systems, positioning them as essential players in both normal physiology and stress-induced responses (Herman et al., 2016).

The synthesis of GCs occurs through the adrenal cortex, where cholesterol serves as precursor molecule (Chakraborty et al., 2021; Marchi and van Eeden, 2021; Miller, 2009). The hormone cortisol (also known as hydrocortisone) is the primary GC produced in humans, whereas corticosterone is more dominant in rodents. Upon stimulation by adrenocorticotropic hormone (ACTH) released by the pituitary gland, the adrenal glands synthesize and secrete GCs, which then circulate in the bloodstream, exerting systemic effects (Kageyama et al., 2021). This tightly controlled synthesis is crucial for physiological balance; dysregulation of GC levels, either by overproduction or deficiency, can have significant effects on health.

Overproduction of GCs can lead to conditions such as Cushing’s syndrome, characterized by symptoms including weight gain, hypertension, hyperglycemia, and immune suppression (Reincke and Fleseriu, 2023). Conversely, an absence or severe deficiency of GCs, such as in Addison’s disease, can result in symptoms such as fatigue, weight loss, hypotension, and potentially life-threatening adrenal crises (Barthel et al., 2019). Both conditions highlight the necessity of maintaining appropriate GC levels for physiological function.

To manage GC dysregulation, synthetic GCs are often administered to either supplement or replace endogenous hormone production. These synthetic compounds are highly potent, with varying degrees of efficacy compared to cortisol (Meikle and Tyler, 1977; Scherholz et al., 2019). Dexamethasone is considered approximately 100-fold more potent than hydrocortisone, while Prednisolone has about 10-fold greater potency compared to hydrocortisone (Meikle and Tyler, 1977). These synthetic GCs are commonly used in clinical settings for their powerful anti-inflammatory and immunosuppressive effects, particularly in the treatment of autoimmune diseases, allergies, and inflammation (Dubois-Camacho et al., 2017; Holgate and Polosa, 2006; Niewoehner et al., 1999; Paolino et al., 2017; Strehl et al., 2019).

GCs exert their effects primarily through binding to the glucocorticoid receptor (GR), a ligand-activated transcription factor of the nuclear receptor superfamily (Cain and Cidlowski, 2017). Upon binding, the GC-GR complex undergoes a conformational change that allows it to translocate into the cell nucleus, where it acts as a transcription factor (Wikstrom et al., 2002). This complex interacts with specific glucocorticoid response elements (GREs) on DNA, leading to either activation or repression of target gene transcription (Hayashi et al., 2004; Lonard and O’Malley B, 2007; Strahle et al., 1987). Through this mechanism, GCs regulate the expression of numerous genes involved in immune modulation, metabolism and cellular stress responses (Weikum et al., 2017). The GR can also interact with other transcription factors, such as NF-κB and AP-1, to inhibit their activity and thereby exert anti-inflammatory effects (Biddie et al., 2011; Kerppola et al., 1993; Ratman et al., 2013; Vettorazzi et al., 2022).

The immune system is a major target of GCs action, and their effects on lymphoid cells, including B lymphocytes, are particularly significant. The GR is expressed in various immune cell types, including B cells, where it mediates the direct effects of GCs on these lymphocytes (Cain and Cidlowski, 2017). GCs have been shown to influence B cell development, differentiation, and function, which includes promoting apoptosis of immature B cells, thereby regulating B cell homeostasis and preventing excessive immune reactions (Gruver-Yates et al., 2014).

GCs play a complex role in immune regulation, particularly in B and T lymphocytes. For instance, the GR is expressed in all B-cell developmental stages and GR-mediated signaling can induce apoptosis in these cells (Gruver-Yates et al., 2014). However, although GCs were shown to induce apoptosis in B and T cells (Gruver-Yates et al., 2014; Herold et al., 2006), some studies suggested that GCs enhance survival in certain contexts (Barczyk et al., 2010; Franchimont, 2004; Vacchio et al., 1998).

Thus, despite the progress in elucidating the role of GCs and their receptor in B cell biology, several questions remain unanswered. Specifically, it remains unclear how different levels of GR activity influence B cell populations and responses under various physiological and pathological conditions. In this study, we addressed the effect of increased GC activity or absent GR expression on the survival of different B cell populations. By addressing the role of BCR expression and B cell activation, we found that IgM BCR expression combined with B cell activation provides relative protection against GC-mediated cell death. Addressing these questions is crucial, as it could provide novel insights into the development of therapeutic strategies aimed at modulating B cell function in autoimmune diseases, allergies, and other immune-related conditions.

## Results

### B cell-specific GR deletion alters B cell subpopulations

To explore the role of the glucocorticoid receptor (GR) in B cell development, we generated B cell-specific GR-deficient mice by crossing GR floxed mice (*GR^f/f^*) with mb1-cre mice, resulting in cre-mediated gene inactivation at the pro-B cell stage. The efficiency of GR deletion was confirmed by flow cytometry, showing significantly reduced GR expression in splenic B cells from *GR^f/f^* x *mb1-cre^+/ki^* mice compared to non-B cells and wild-type (WT) controls (Fig. S1A). We observed an increase in the frequency and absolute numbers of small CD25^+^ pre-B cells (Fig. S1B-C), although the deletion did not lead to an overall increase of total B cell numbers in the bone marrow (Fig. S1C).

In the spleen, *GR^f/f^* x *mb1-cre^+/ki^*mice displayed a higher percentage and absolute number of total B cells (Fig. 1A). Further analysis of splenic B cell subpopulations revealed significant increases in the absolute numbers of follicular B cells (Fo.B) and CD21^low^ (CD21^lo^.B) B cells, despite unchanged percentages (Fig. 1B-C). Interestingly, the percentage of marginal zone B cells (MZ.B) was significantly reduced, although their absolute numbers remained unaffected (Fig. 1B-C).

**Figure 1.**
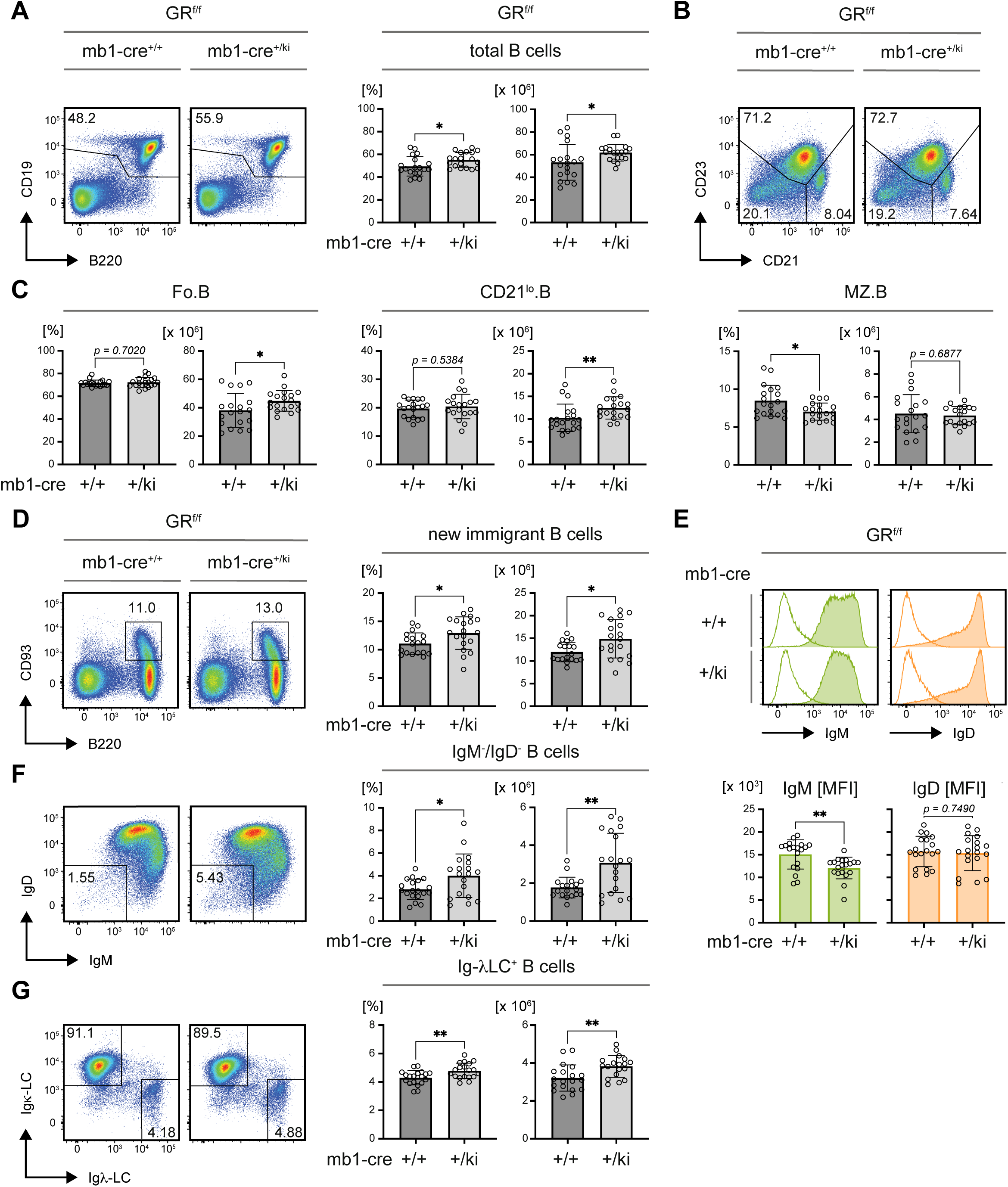
B cell-specific GR deletion alters splenic B cell subpopulations. Phenotype analyses of B cell populations in 8 weeks old glucocorticoid receptor (GR) *GR^f/f^* x *mb1-cre* mice of the indicated genotype, n = 19 each group; mean ± SD. If not indicated otherwise, flow cytometric data were pre-gated on B cells as shown in **A** (left panel). Statistical significance was calculated by applying either the unpaired t test or Mann-Whitney-U test depending on the distribution of data points. **A** Representative flow cytometric analysis of total splenic B cells (left panel) pre-gated on single viable cells and quantification of percentages (bar diagrams, left) and absolute cell numbers (bar diagrams, right). **B** Representative flow cytometric analysis of follicular (Fo.B: CD23^+^/CD21^+^) marginal zone (MZ.B: CD23^-^/CD21^+^) and CD21^lo^ (CD21^lo^.B: CD23^-^/CD21^lo^) B cells in the spleens of mice from the indicated genotypes. **C** Quantified percentages and absolute numbers of Fo.B cells (left), CD21^lo^.B (middle) and MZ.B cells (right) in the spleens of *GR^f/f^* x *mb1-cre^+/+^* and *GR^f/f^* x *mb1-cre^+/ki^* mice. **D** Representative flow cytometric analysis of new immigrant B cells in spleens (left panel) and quantification of percentages (bar diagrams, left) and absolute cell numbers (bar diagrams, right). *GR^f/f^* x *mb1-cre^+/+^*, n = 18. **E** Representative flow cytometric analysis of IgM (green) and IgD (orange) surface expression in total splenic B cells (top panel) and quantification of the respective mean fluorescence intensities (MFI; bottom panel). **F** Representative flow cytometric analysis of IgM^-^/IgD^-^ splenic B cells (left panel) and quantification of percentages (bar diagrams, left) and absolute cell numbers (bar diagrams, right). **G** Representative flow cytometric analysis of intracellular Ig-κ & −λLC expression in splenic B cells (left panel) and quantified percentages of Ig-λLC^+^ B cells (bar diagrams, left) and absolute cell numbers (bar diagrams, right).

The CD21^lo^ B cell compartment, which includes transitional B cells and B-1 cells, was expanded, and the population of new immigrant B cells was also elevated in GR-deficient mice (Fig. 1D). Notably, these cells exhibited reduced surface expression of IgM, while IgD expression remained unaffected (Fig. 1E). Additionally, an increased population of IgM/IgD double-negative B cells was observed in the spleens of GR-deficient mice (Fig. 1F), accompanied by a significant accumulation of B cells expressing the lambda light chain (Ig-λLC), while kappa light chain (Ig-κLC) expressing cells remained unchanged (Fig. S1D).

Further characterization of the IgM^-^/IgD^-^ population showed that these cells were predominantly located in the CD21^lo^ B cell compartment (Fig. S1E). This population likely includes plasma cells, since ELISpot assays detected IgM- and IgG-secreting cells primarily within the CD21^lo^ B cell population (Fig. S1F). Interestingly, the IgM^-^/IgD^-^ B cell population also contained a fraction of isotype-switched IgG^+^ memory B cells (Fig. S1G).

However, IgM^-^/IgD^-^ B cells from GR-deficient mice did not show a significant increase in their ability to secrete IgM in ELISpot (Fig. S1H), and the overall concentration of serum IgM and IgG remained unchanged (Fig. S1I). Moreover, no significant increase in CD138^+^ plasma cells was detected in GR-deficient mice (Fig. S1J).

In conclusion, deletion of the GR in B cells leads to changes in specific B cell subsets, including an increase in follicular and CD21^low^ B cells, and an accumulation of IgM^-^/IgD^-^ B cells. These results suggest that the GR plays a role in regulating B cell homeostasis, possibly by eliminating certain B cell subsets that would otherwise persist in the absence of GR-mediated signaling.

### Continuous glucocorticoid treatment eradicates B cells *in vivo*

To investigate the effects of sustained GR activation on B cells, we treated mice with the synthetic GR agonists, Dexamethasone (Dex) and Prednisolone (Pred), using continuous-release pellets over a period of 14 days. Since Dex is approximately ten times more potent than Pred due to differences in stability, half-life, and bioavailability, Pred was administered at a ten-fold higher concentration to attempt comparable effects *in vivo* (Meikle and Tyler, 1977).

Both Dex and Pred significantly reduced the percentage of total B cells in the bone marrow (Fig. S2A) and total splenocyte numbers compared to sham-treated controls (Fig. 2A). However, Pred treatment was more effective, as shown by a significant reduction in spleen weight (Fig. 2B) and absolute B cell numbers in the bone marrow, which was not be observed upon Dex treatment (Fig. S2A). Additionally, both treatments led to a significant suppression of endogenous corticosterone levels, suggesting interference with the body’s natural glucocorticoid biosynthesis.

**Figure 2.**
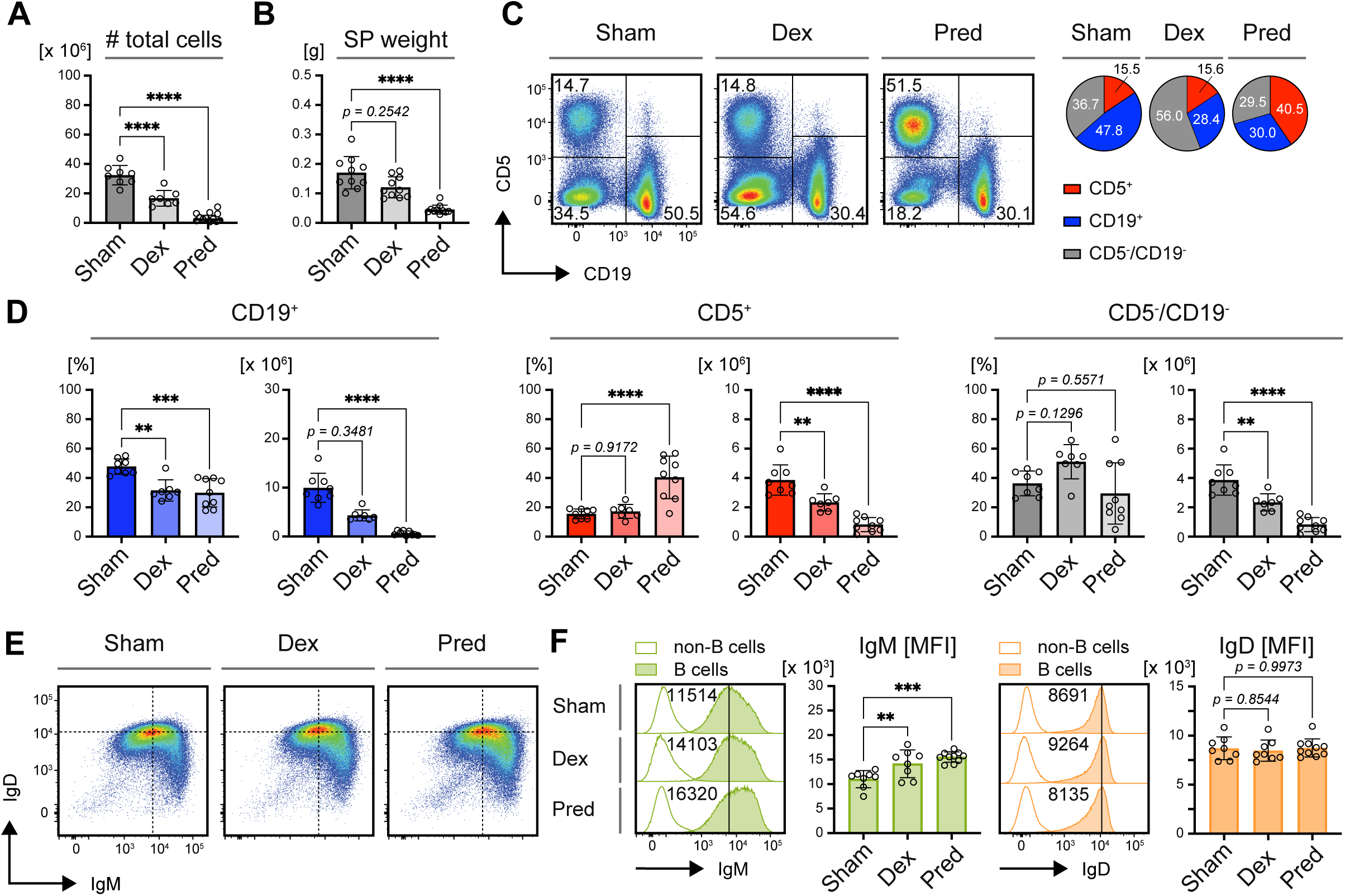
Continuous GC-treatment eradicates B cells *in vivo*. Phenotype analyses of mice transplanted with constant glucocorticoid (GC)-release pellets after 14 days of GC treatment. **A** Total cell numbers in spleens from mice following 14 days of exposure to Dexamethasone (Dex, n = 7), Prednisolone (Pred, n = 8) or control (Sham, n = 10) pellets. Mean ± SD. Statistical significance was calculated by applying the Kruskal-Wallis test. **B** Quantified spleen weight of mice after 14 days of exposure to Dex (n = 10), Pred (n = 12) or Sham (n = 10) pellets. Mean ± SD. Statistical significance was calculated by applying the Kruskal-Wallis test. **C** Representative composition of residual cells in spleens after 14 days of treatment with the indicated pellets as measured by flow cytometry (left panel). Cells were pre-gated on single viable lymphocytes. Pie charts in the right panel show average percentages of B cells (CD19^+^), T cells (CD5^+^) & non-B/T cells (CD5^-^/CD19^-^), representative of at least 7 individual mice per group. **D** Quantification of percentages (left panels) and absolute cell numbers (right panels) of B cells (CD19^+^, blue), T cells (CD5^+^, red) & non-B/T cells (CD5^-^/CD19^-^, gray) in murine spleens after 14 days of treatment with Sham (n = 8), Dex (n = 7) and Pred (n = 9) pellets. Mean ± SD. Statistical significance was calculated by applying the ordinary one-way ANOVA or the Kruskal-Wallis test, respectively. **E** Representative flow cytometric analysis of IgM and IgD surface expression in total splenic B cells (pre-gated on single viable lymphocytes) purified from mice of the indicated treatment cohorts. **F** Representative comparison of IgM (green, left) and IgD (orange, right) MFI in total splenic B cells (pre-gated on single viable lymphocytes) from mice of the indicated treatment groups (histograms). Non-B cells within the respective samples served as negative controls. Bar diagrams show quantification of IgM and IgD MFI in Sham- (n = 8), Dex- (n = 7) and Pred-treated (n = 9) mice. Mean ± SD. Statistical significance was calculated by applying the Kruskal-Wallis test.

To assess the remaining lymphocyte populations after 14 days of treatment, splenocytes were stained with anti-CD5 and -CD19 antibodies (Fig. 2C). The frequency of B cells, which typically represent about half of the splenic cells in sham-treated controls, was drastically reduced to less than one-third of the remaining cells. Interestingly, although the total numbers of T cells were reduced, their frequency in the residual splenic cell population remained constant or even increased following Pred treatment (Fig. 2D). Non-B/non-T cells also showed a trend towards an increased frequency in Dex-treated mice, indicating that T cells and non-B/non-T cells were more resistant to glucocorticoid-induced depletion than B cells and that T cells tolerate a high concentration of Pred.

Given the previously observed reduction in IgM expression following GR inactivation (Fig. 1E), we analyzed IgM and IgD expression on B cells that survived glucocorticoid treatment. The IgM/IgD profile resembled that of sham-treated controls (Fig. 2E), but a notable shift towards higher IgM expression was observed in GR agonist-treated mice. Flow cytometric analysis revealed that IgM expression was significantly upregulated following treatment with GR agonists (Fig. 2F), contrasting with the decreased IgM levels seen in GR-deficient B cells. Although IgD expression remained unchanged, the increase in surface IgM significantly raised the IgM/IgD ratio (Fig. S2B).

In conclusion, these results indicate that continuous GC treatment selectively promotes the survival of IgM^high^ B cells, suggesting that these cells are favored under GC exposure, potentially as a mechanism to escape GC-mediated cell death. Additionally, B cells appear to be highly sensitive to GC-induced depletion compared to other immune cells such as T cells and non-lymphoid/myeloid cells, which are less affected under the same treatment conditions.

### Activated B cells display relative resistance to treatment with GR agonists

To further investigate the effects of glucocorticoid receptor (GR) activation on B cells, we purified mature splenic B cells from WT mice and treated them *in vitro* with the GR agonists Dex or Pred (Fig. 3A). Even at a low nanomolar concentration (25 nM), GR agonist exposure led to the complete eradication of all viable B cells within 3 days (Fig. 3B). In contrast, mature B cells from *GR^f/f^*x *mb1-cre^+/ki^* mice, which lack functional GRs in B cells, were resistant to the treatment (Fig. S3A), confirming that the cell loss was specifically mediated by GR activation.

**Figure 3.**
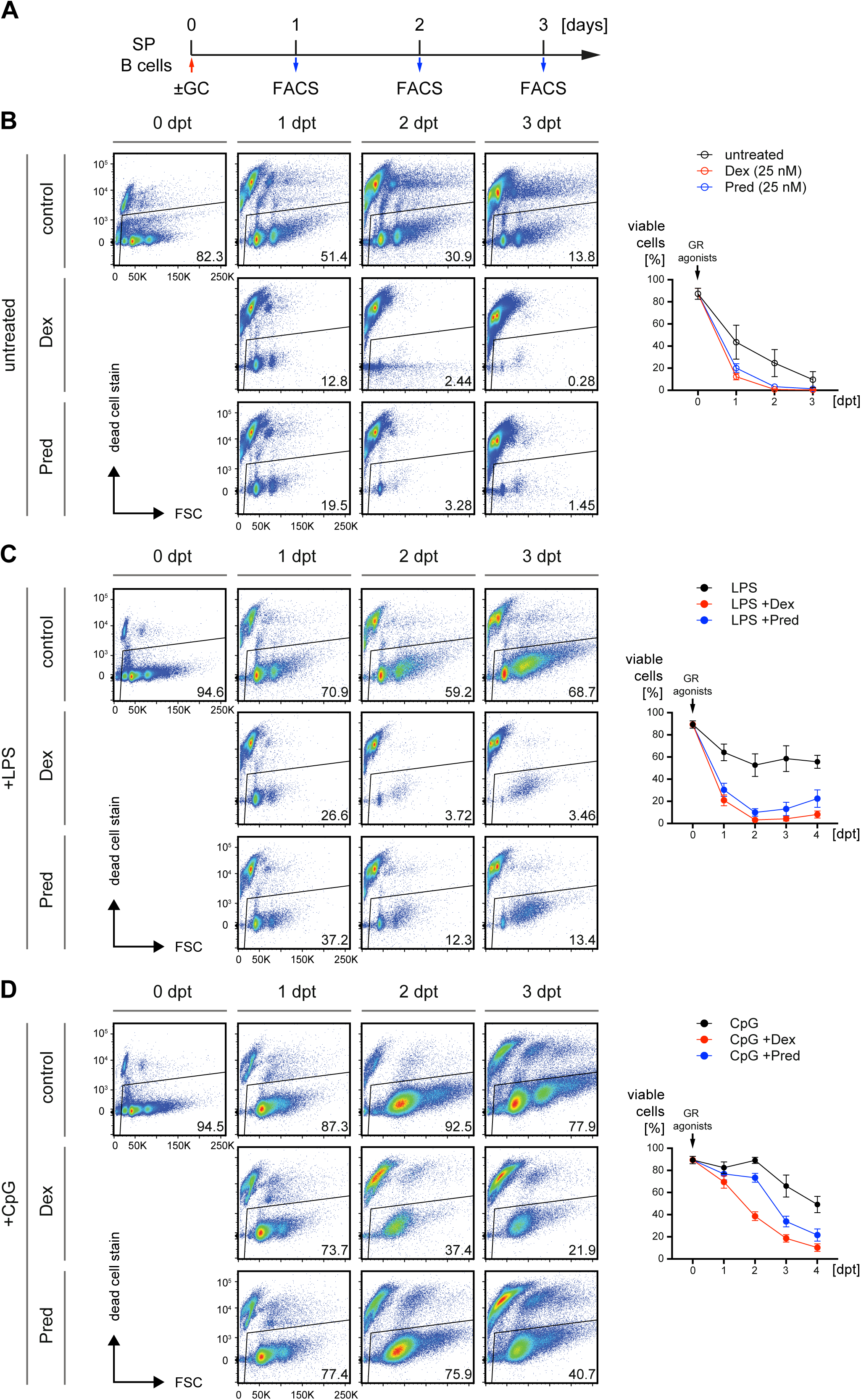
Activated mature B cells are more resistant to GR agonist treatment *in vitro*. **A** Schematic timeline of the experimental procedure: Mature splenic B cells from wild-type (WT) mice were purified by negative selection via magnetic activated cell sorting (MACS) and treated with the indicated GR agonists at a concentration of 25 nM. Cell viability was monitored by flow cytometry on days 1 to 4 post treatment (dpt). **B** Representative flow cytometric analyses of cell viability upon *in vitro* treatment with GR agonists (left panel) as described in **A** and quantified percentages of viable cells. Control (black, n = 22), Dex (red, n = 14), Pred (blue, n = 14), mean ± SD. **C** Representative flow cytometric analyses of cell viability upon simultaneous treatment with 2.5 µg/mL lipopolysaccharide (LPS) and GR agonists (left panel) *in vitro* as described in **A** and quantified percentages of viable cells. n = 17 in each group, mean ± SD. **D** Representative flow cytometric analyses of cell viability upon concomitant treatment with 2 µM CpG-ODN #1826 and GR agonists (left panel) *in vitro* as described in **A** and quantified percentages of viable cells. n = 17 in each group, mean ± SD.

To address the effect of cell activation on GC-mediated cell death, we used the toll-like receptor 4 (TLR4) agonist lipopolysaccharide (LPS) to simultaneously activate B cells during GR agonist treatment (Fig. 3C). LPS activation of B cells led to a minor improvement in survival compared to resting cells, with a small proportion surviving until day 3 of treatment. Since TLR9 activation by unmethylated CpG oligonucleotides (CpG) typically induces stronger and faster activation than LPS, we also tested whether CpG could better protect B cells from GR agonist-mediated cell death (Fig. 3D). Remarkably, co-stimulation with CpG significantly rescued B cells from death when applied simultaneously with GCs, further demonstrating that activated B cells are more resistant to GR-induced cell death than resting B cells.

These findings suggest that although both LPS and CpG act through MyD88- dependent pathways (O’Neill and Bowie, 2007), the activation profiles they induce may differ in strength, speed, and their capacity to rescue B cells from GC-induced apoptosis. Moreover, resting B cells isolated from the spleen and cultured *in vitro* in the presence of GR agonists are highly sensitive to GC-mediated cell death, but activation via TLRs significantly improves survival.

### GR agonists enhance B cell activation and accelerate terminal differentiation

The data above allow the conclusion that CpG provides more effective protection through rapidly and robustly activating B cells, thereby enabling them to withstand GC-induced apoptosis. To confirm this conclusion, we analyzed the expression of the activation markers CD69 and CD86 on WT B cells treated with either TLR ligand (Fig. S4A). CpG treatment led to rapid and nearly complete upregulation of CD69, with >90% of cells showing elevated expression one day post-treatment (dpt). In contrast, LPS treatment resulted in elevated CD69 expression in <50% of cells. Interestingly, while CD86 expression remained low and declined in CpG-treated cells over time, LPS-treated B cells exhibited a marked upregulation of CD86, which increased steadily until 3 dpt (Fig. S4A).

When combined with GR agonist treatment (Dex or Pred), both LPS- and CpG-activated B cells showed a significant enhancement in the expression of CD69 and CD86 (Fig. 4A-B). Interestingly, treatment of splenic B cells with GR agonists in absence of LPS or CpG, already resulted in up-regulation of the activation marker CD69 at 1 and 2 dpt, whereas CD86 was only upregulated at day 2 in the few residual cells (Fig. S4B). Following concomitant treatment with LPS, CD69 expression was significantly increased upon addition of Dex or Pred while CD86 was decreased following Dex or Pred treatment on day 1 and later increased on 3 dpt (Fig. 4A). On the other hand, CpG-treated B cells exhibited increased expression of both CD69 and CD86, which was not further increased at 1 day after Dex or Pred treatment (Fig. 4B). However, on the following days, Dex or Pred treatment led to an increase in CD86 expression as compared with cells treated only with CpG. Together, these data suggest that rapid and sustained CD69 upregulation distinguishes CpG from LPS induced splenic B cells and that CG treatment selects these cells.

**Figure 4.**
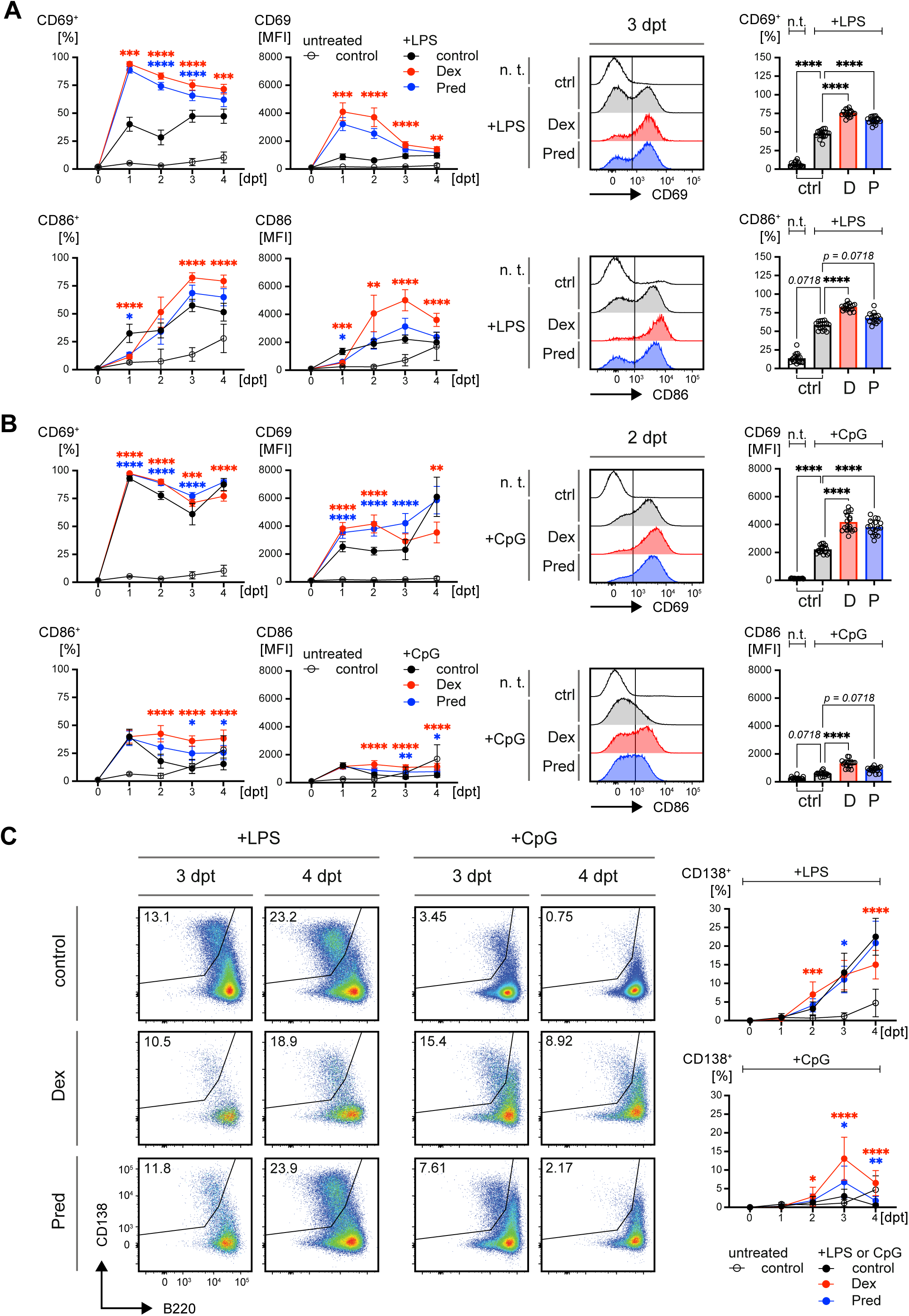
GR agonists enhance B cell activation and accelerate terminal differentiation. **A - B** Kinetics of activation markers CD69 (top panel) and CD86 (bottom panel), determined by flow cytometry in WT splenic B cells following treatment with GR agonists as described in Fig. 3A in presence of LPS (**A**) or CpG (**B**). Representative flow cytometric analyses and bar diagrams on the right-hand side compare percentages of CD69 and CD86 positive cells upon LPS-treatment at 3 dpt (**A**) or the respective MFIs for CpG-treatment at 2 dpt (**B**). MFIs were used because, in contrast to LPS, all CpG treated cells showed immediate upregulation of CD69. For all groups and time points, n = 17 except for 0 dpt (n = 11) and 1 dpt (n = 14), mean ± SD. Statistical significance was calculated by applying either the RM one-way ANOVA or the Friedman test, respectively. Asterisks in the kinetics indicate significant differences between cells stimulated either with LPS or CpG and cells additionally treated with GR agonists Dex (D, red asterisks) or Pred (P, blue asterisks). **C** Representative flow cytometric analysis of *in vitro* plasma cell differentiation at days 3 and 4 following treatment of WT splenic B cells with GCs in presence of either LPS (left) or CpG (right). The changes of CD138^+^/B220^lo^ cells over time are shown as kinetics on the right-hand side. For all groups and time points, n = 17, mean ± SD. Statistical significance was calculated by applying either the RM one-way ANOVA or the Friedman test, respectively. Asterisks in the kinetics indicate significant differences between cells stimulated either with LPS or CpG and cells additionally treated with GR agonists Dex (D, red asterisks) or Pred (P, blue asterisks).

Next, we explored whether addition of GR agonists influences plasma cell differentiation, which is typically induced by LPS *in vitro*. While Dex or Pred treatment did not further enhance the plasma cell differentiation capacity of LPS-stimulated B cells (Fig. 4C), an interesting observation was made in CpG-treated cells: Although CpG alone was insufficient to induce efficient plasma cell differentiation, the addition of GR agonists led to the generation of CD138^+^/B220^lo^ plasma cells (Fig. 4C).

Together, the data suggest that CpG and LPS differ in their kinetics of B cell activation and that GR agonists amplify this activation. Moreover, the GR agonists boosted plasma cell differentiation of B cells treated with CpG, which was not observed for LPS.

### Delayed GR agonist treatment after LPS activation improves B cell survival

The data presented above suggested that B cell activation plays a crucial role in protecting cells from GC-induced cell death. Based on CD69 expression, CpG was shown to induce faster B cell activation than LPS, likely explaining its superior ability to protect B cells during simultaneous treatment with GR agonists. Therefore, we hypothesized that LPS might offer protection if GR agonist treatment is delayed, allowing sufficient time for activation to occur.

To directly test this, splenic B cells were pre-incubated with LPS, and GR agonists were added two days post LPS treatment, with cells analyzed in subsequent days (Fig. 5A). The delayed GR agonist treatment resulted in substantial rescue of the LPS-activated B cells at 3 dpt (Fig. 5B, Fig. S5), compared to simultaneous treatment with LPS and GR agonists (Fig. 5C). Moreover, delayed treatment of LPS-stimulated cells with GR agonists led to a significant increase in the proportion of plasma cells observed on days 3 and 4 (Fig. 5D-E). Notably, GR-deficient splenic B cells were also resistant to delayed treatment with either Dex or Pred (Fig. S6A) and did not show the enhanced plasma cell differentiation as compared with cells from control mice (Fig. S6B).

**Figure 5.**
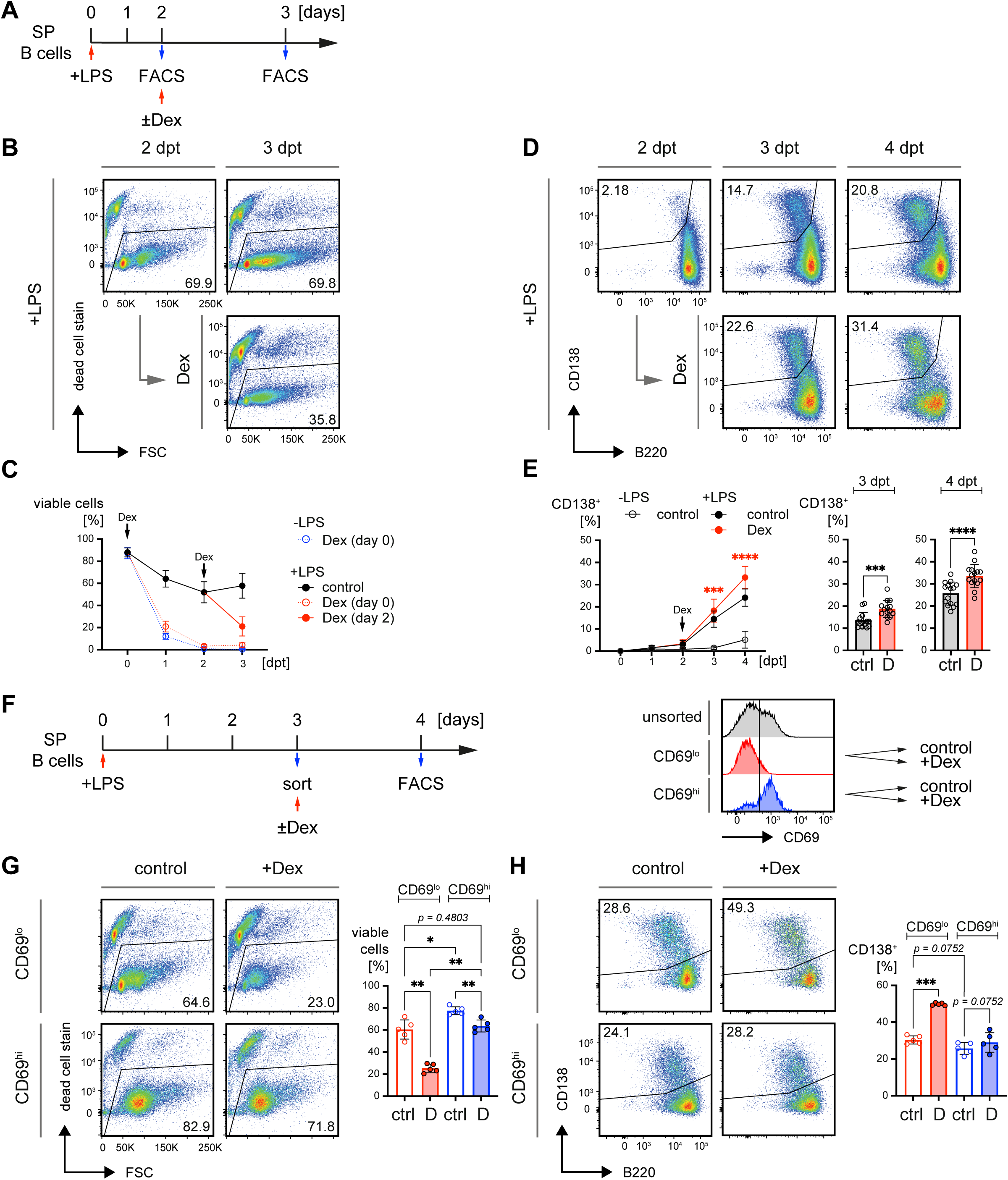
Delayed GR agonist treatment after LPS activation improves B cell survival. **A** Schematic timeline of the experimental procedure: Mature splenic B cells from WT mice were isolated by negative selection via MACS and treated with 2.5 µg/mL LPS. On day 2, 25 nM Dex (D) were added or the cells were left untreated (ctrl). Cell viability was monitored by flow cytometry one day later (3 dpt). **B** Representative flow cytometric analyses of cell viability at day 2 and 3 in LPS pre-stimulated B cells upon addition of Dex at day 2. **C** Kinetics comparing survival of mature B cells when treated with Dex at day 0 (dotted lines, data already shown in Fig. 3B) in presence (red) or absence of LPS (blue) to survival, when Dex was added at day 2 (solid lines and filled symbols), n ≥ 14, mean ± SD. **D** Representative flow cytometric analyses of plasma cell differentiation at the indicated time points in LPS pre-stimulated B cells upon delayed addition of 25 nM Dex, as described in **A**. Day 0 & 1: n = 11; days 2 - 4: n = 15; mean ± SD. **E** Kinetics in the left panel show quantified percentages of CD138^+^/B220^lo^ cells when Dex (D) was added on day 2 (n = 6) upon pre-stimulation with 2.5 µg/mL LPS. Bar diagrams on the right-hand side compare percentages of CD138^+^/B220^lo^ cells in the indicated treatment groups at 3 and 4 dpt. Mean ± SD. Statistical significance was calculated by applying the paired t test for day 2 and either the RM one-way ANOVA or the Friedman test for assessing differences at days 3 and 4. Asterisks in the kinetics indicate significant differences between cells stimulated with LPS and cells additionally treated with GR agonists Dex (D, red asterisks). **F** Schematic timeline of the experimental procedure: Mature splenic B cells from WT mice were purified by negative selection via MACS and treated with 2.5 µg/mL LPS. On day 3 CD69^lo^ and CD69^hi^ B cells were FACS-purified according to the gating strategy shown in **Fig. S4C** and treated overnight in presence (D) or absence (ctrl) of 25 nM Dex, n = 5, mean ± SD. Cell viability was monitored by flow cytometry one day later (4 dpt). **G** Representative flow cytometric analyses of cell viability in FACS-purified CD69^lo^ and CD69^hi^ B cells upon treatment with Dex (D, left panel) as described in **F** and quantified percentages of viable cells, n = 5 in each group, mean ± SD. Statistical significance was calculated by applying the RM one-way ANOVA. **H** Representative flow cytometric analyses of *in vitro* plasma cell differentiation in FACS-purified CD69^lo^ and CD69^hi^ B cells upon treatment with Dex (D, left panel) as described in **A** and quantified percentages of CD138^+^/B220^lo^ cells, n = 5 in each group, mean ± SD. Statistical significance was calculated by applying the RM one-way ANOVA.

To determine whether LPS-induced activation was responsible for the improved survival of B cells, we sorted LPS-activated B cells based on CD69 expression on day 3 (Fig. S7A; Fig. 5F). Time-course experiments had previously revealed that CD69 expression peaked on this day (Fig. S4A). Subsequent addition of Dex to those sorted cells further increased CD69 expression in CD69^low^ and CD69^high^ populations, whereas CD86 levels remained unchanged (Fig. S7B). Moreover, GR agonist treatment demonstrated that CD69^high^ B cells were significantly more resistant to GC-mediated cell death compared to CD69^low^ cells (Fig. 5G). Remarkably, Dex-treated CD69^high^ cells exhibited survival rates similar to untreated CD69^low^ cells, suggesting that activation confers CD69^high^ cells a relative resistance to GC-induced cell death.

Interestingly, when examining plasma cell differentiation, GR agonist treatment induced a significant increase in plasma cells in the CD69^low^ population but not in the CD69^high^ population (Fig. 5H). Together, these data suggest that the induction of activation markers, such as CD69, is associated with protection from GC-mediated cell death, while cells lacking activation markers show an increased propensity for plasma cell differentiation upon GR agonist treatment.

### IgD BCR expression affects resistance to cell death

Similar to the regulation of activation markers like CD69 and CD86, CpG treatment led to distinct kinetics in the modulation of IgM/IgD BCR expression on splenic B cells *ex vivo*. Specifically, CpG induced a rapid downregulation of IgD in the entire cell population, whereas LPS caused a delayed yet sustained downregulation of IgD only in a subfraction of the *in vitro*/*ex vivo* cultured splenic B cells (Fig. S8A). This difference in BCR expression mirrored the kinetics observed with activation markers and seemed to impact how B cells responded to GR agonist treatment (Fig. S4A).

When we treated the LPS-stimulated cells with GR agonists, IgD^high^ B cells proliferated less efficiently and were particularly susceptible to GC-induced cell death (Fig. 6A) in a GR-dependent manner (Fig. S8B). In contrast, IgD^low^ (IgM^high^) cells, which increased following LPS treatment, were further enriched when exposed to GR agonists. This pattern suggested that IgD downregulation, particularly in IgM^high^ B cells, was linked to increased survival under GR agonist treatment (Fig. 6A).

**Figure 6.**
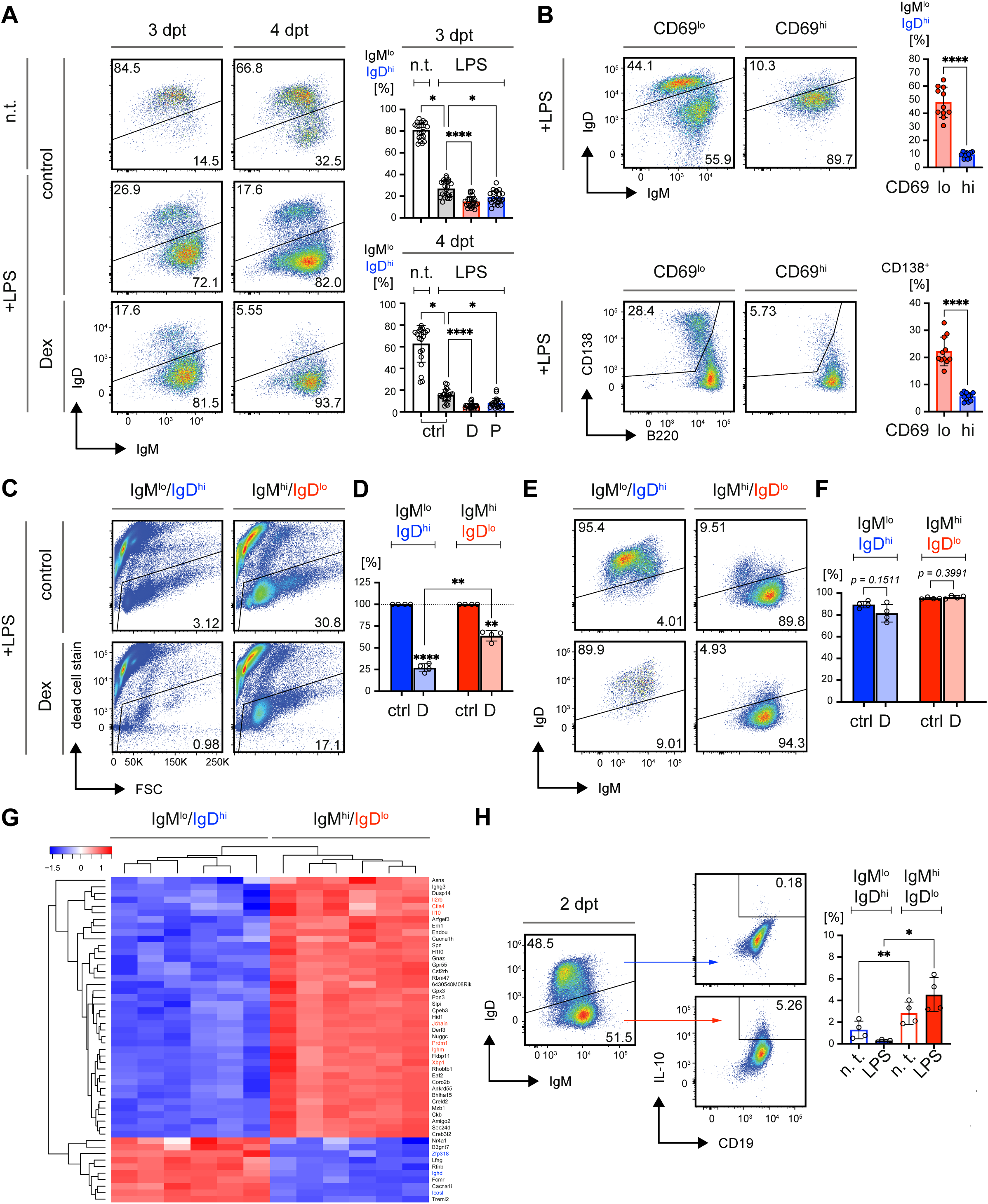
IgD BCR expression affects resistance to cell death. **A** Representative flow cytometric analyses of IgM and IgD surface expression in mature splenic B cells upon stimulation with LPS and delayed treatment with GR agonists as described in Fig. 5D. Bar diagrams below depict quantified percentages of B cells with IgM^lo^/IgD^hi^ surface expression at 3 and 4 dpt. For day 3: n = 22; day 4: n = 21; mean ± SD. Statistical significance was calculated by applying the Friedman test. **B** Representative flow cytometric analyses of IgM/IgD surface expression (top) and plasma cell differentiation (CD138/B220 bottom) in CD69^hi^ and CD69^lo^ splenic WT B cells at day 3 upon stimulation with 2.5 µg/mL LPS. Bar diagrams besides the flow cytometry data show quantified percentages of IgD^hi^ cells and CD138^+^ cells in the indicated cell populations. n = 11; mean ± SD. Statistical significance was calculated by applying the paired t test. **C** Representative flow cytometric analysis of viability in FACS-purified IgM^lo^/IgD^hi^ and IgM^hi^/IgD^lo^ mature splenic B cells at 3 dpt upon sorting and treatment with Dex as indicated in the schematic timeline displayed in Fig. 5F. Mature splenic B cells from WT mice were purified by negative selection via MACS and stimulated with 2.5 µg/mL LPS. After 2 days of incubation, cells were FACS-purified to separate IgM^lo^/IgD^hi^ and IgM^hi^/IgD^lo^ populations according to the gating strategy in **Fig. S9D**. Subsequently Dex was added to the culture at a concentration of 25 nM. Viability and B cell receptor (BCR) expression was monitored by flow cytometry at 3 dpt. **D** Quantified percentages of viable cells from **D** normalized to viability in LPS-treated cells and shown as fold change (%). n = 4, mean ± SD. Statistical significance between control- and Dexamethasone (D)-treated cells was calculated by applying the one sample t test, while differences between D-treated IgM^lo^/IgD^hi^ and IgM^hi^/IgD^lo^ were assessed by using the RM one-way ANOVA. **E** Representative flow cytometric analysis of IgM and IgD surface expression in purified mature splenic B cells at 3 dpt upon sorting and Dex treatment as described in **D**. **F** Quantified percentages of IgM^lo^/IgD^hi^ (blue) and IgM^hi^/IgD^lo^ (red) in purified B cells from **F**. n = 4, mean ± SD. Statistical significance between control- and Dexamethasone (D)-treated cells was calculated by applying the RM one-way ANOVA. **G** Heatmap of the 50 most up- (red) or down-regulated (blue) transcripts in IgM^lo^/IgD^hi^ and IgM^hi^/IgD^lo^ splenic B cells from WT mice as determined by RNA deep sequencing. Mature splenic B cells from WT mice were purified by negative selection via MACS and stimulated with 2.5 µg/mL LPS. After 3 days of incubation, cells were FACS- purified to separate IgM^lo^/IgD^hi^ and IgM^hi^/IgD^lo^ populations for transcriptome analysis. Shown data are representative of IgM^lo^/IgD^hi^ and IgM^hi^/IgD^lo^ samples from 6 individual mice and also enlisted in **Table S1**. **H** Representative flow cytometric analysis of IL-10 expression and quantification of IL-10^+^ percentages at 4 dpt in IgM^lo^/IgD^hi^ and IgM^hi^/IgD^lo^ splenic WT B cells upon cultivation in presence or absence (n. t., **Fig. S10C**) of 2.5 µg/mL LPS. N = 4, mean ± SD. Statistical significance between untreated- and LPS-stimulated cells was calculated by applying the RM one-way ANOVA.

Importantly, IgD^low^ (IgM^high^) expression correlated with higher CD69 expression and decreased plasma cell differentiation (Fig. 6B), suggesting that the enhanced survival of CD69^high^ B cells might be partially caused by reduced IgD BCR expression. In fact, IgD^high^ cells showed increased cell death on day 3 and 4 after Dex treatment (Fig. 6A). To test whether IgD downregulation was critical for protection from GC-induced cell death, we examined the survival of sorted IgD^high^ (IgM^low^) and IgD^low^ (IgM^high^) splenic B cells on day 2 after LPS treatment *in vitro* (Fig. S9). These sorted populations were then subjected to overnight Dex treatment. The results revealed significantly higher levels of cell death in IgD^high^ (IgM^low^) cells compared to IgD^low^ (IgM^high^) cells, suggesting that the IgD/IgM BCR expression ratio plays a crucial role in determining susceptibility to glucocorticoid-induced cell death (Fig. 6C-D).

To investigate whether Dex directly influences IgM/IgD BCR expression or merely induces enrichment of the IgD^low^ (IgM^high^) population, we analyzed BCR expression after overnight treatment with Dex. The data showed that cells expressing IgD did not downregulate IgD BCR but were selectively eliminated by the treatment, resulting in an increased proportion of IgD^low^ (IgM^high^) cells (Fig. 6F).

These findings suggest that IgD downregulation in activated B cells is crucial for their protection from GC-induced cell death, highlighting the importance of the IgM^high^/IgD^low^ phenotype in conferring resistance to GC-induced apoptosis.

### IgD^low^ B cells show enhanced survival and increased IL-10 expression

To further understand the differences between IgD^high^ (IgM^low^) and IgD^low^ (IgM^high^) cells, we sorted *in vitro* LPS-activated splenic B cells based on their IgM/IgD expression (Fig. S9) and performed transcriptome analysis through deep sequencing. Comparing the respective subpopulations purified from 6 individual mice, we found that expression of genes associated with plasma cell differentiation, such as *Prdm1*, *J chain*, and *Xbp1*, was significantly upregulated in IgD^low^ (IgM^high^) cells (Fig. 6G, Table S1 positions 10, 2 and 46 respectively). In contrast, genes necessary for IgD expression, including *Zfp318* and *Ighd,* were upregulated in IgD^high^ (IgM^low^) cells (Fig. 6G, Table S1 positions 12 and 5, respectively).

To investigate the mechanism underlying the improved survival of IgD^low^ (IgM^high^) B cells, we analyzed their altered gene expression compared with IgD^high^ (IgM^low^) cells. Although there were no clear differences in pro- or anti-apoptotic genes at the transcriptional level, we observed upregulated expression of genes associated with T cell survival and proliferation, such as *Il2rb* and *CD28*, in the IgD^low^ (IgM^high^) population (Fig. 6G, Table S1 positions 29 and 54, respectively). In line with these findings, KEGG pathway analysis revealed distinct differences in cytokine signaling pathways between the two cell populations (Fig. S10).

To validate these observations, we performed FACS staining with specific antibodies to confirm the gene expression data. Interestingly, IgD^low^ (IgM^high^) cells also exhibited increased expression of IL-10, a cytokine known for its role in regulating immune responses (Fig. 6G, Table S1 position 9). This finding is particularly notable in light of previous research showing that GCs promote the survival of anti-inflammatory macrophages involved in resolving inflammation (Barczyk et al., 2010). Together, our data suggest that GR agonists might selectively enrich for anti-inflammatory or regulatory B cells producing IL-10 or induce IL-10 expression directly. To validate this, we performed intracellular staining for IL-10 on LPS-activated cells and found that IgD^low^ (IgM^high^) cells contained higher amounts of IL-10 (Fig. 6H, Fig. S11A-B). Similar to IL-10, expression of CD122 (encoded by *Il2rb*) was also elevated on protein level in IgD^low^ (IgM^high^) B cells as compared with IgD^high^ (IgM^low^) following LPS stimulation (Fig. S11C). Together with the IL-2 receptor alpha chain (CD25) and the common gamma chain (CD132), CD122 (IL-2 receptor beta chain) forms the IL-2 receptor complex (Malek, 2008).

These results indicate that IgD downregulation is crucial for protecting B cells from GC-induced cell death by activation of cytokine signaling and that GR agonist treatment promotes the enrichment of IgD^low^ (IgM^high^) cells, which may possess regulatory B cell functions based on their IL-10 expression.

### Selective effects of GR agonist treatment on IgD expressing B cells *in vivo*

The previous results suggested that B cells with increased IgD expression are more sensitive to GC-induced cell death, while those with downregulated IgD BCR expression are more resistant. To confirm this finding *in vivo*, we analyzed splenic B cells from mice treated with Dex or Pred release pellets shown above.

After 14 days of GR agonist treatment, we observed a selective reduction in follicular B cells (Fo.B), which was proportional to the potency of the GR agonist used (Fig. 7A). Follicular B cells, known for their higher IgD expression, were particularly sensitive to GR-mediated cell death (Fig. 7B). In contrast, marginal zone (MZ.B) B cells, which typically express lower levels of IgD, were relatively enriched following GR agonist treatment (Fig. 7A-B).

**Figure 7.**
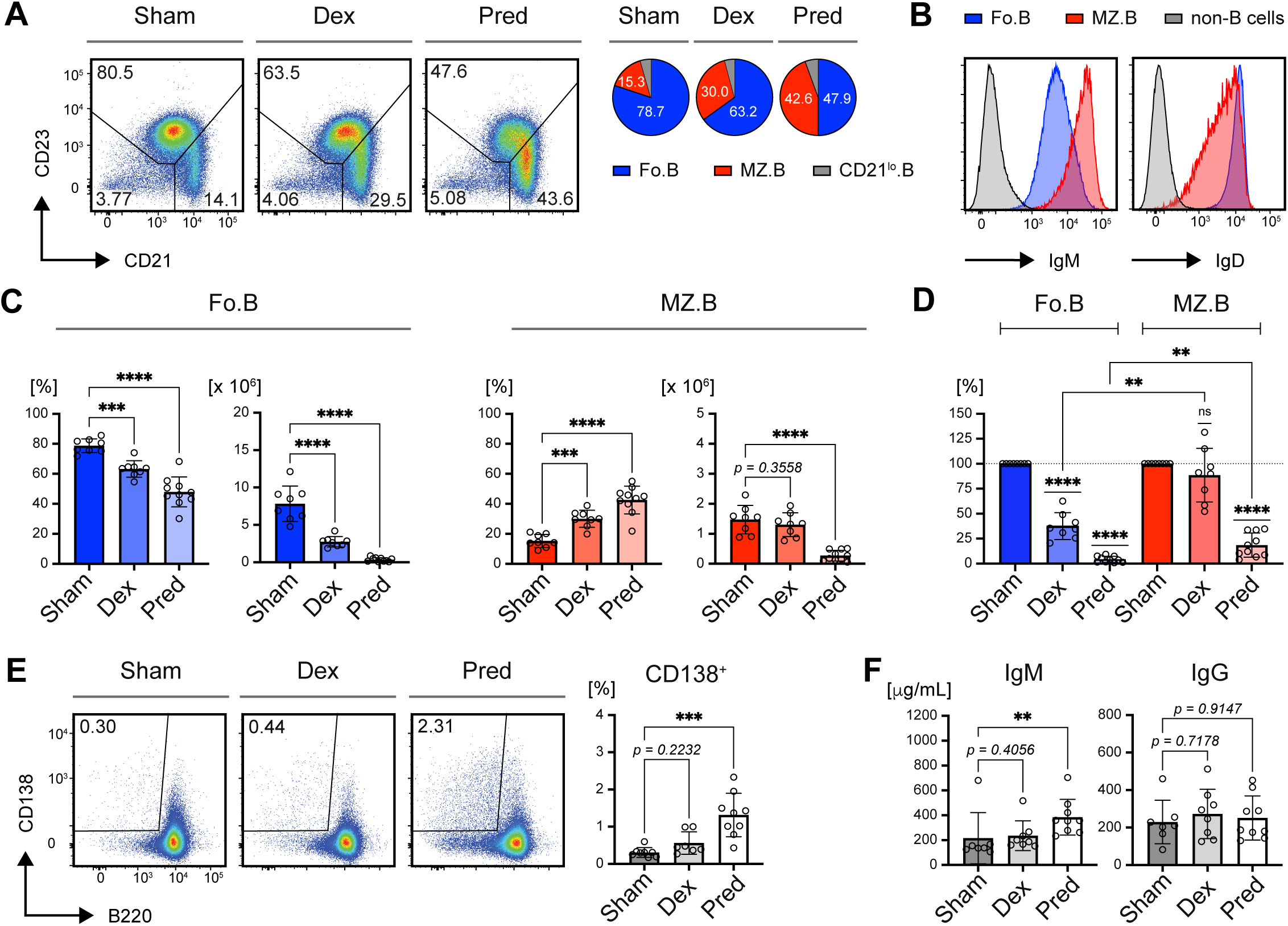
Selective effects of GR agonist treatment on IgD expressing B cells. *in vivo* Phenotype analyses of mice transplanted with constant glucocorticoid (GC)-release pellets after 14 days of GC treatment. **A** Representative flow cytometric analysis of IgM and IgD surface expression in the B cell subpopulations Fo.B, MZ.B and CD21^lo^.B, following GC treatment *in vivo*. Pie charts in the right panel show average percentages of the respective B cell subpopulations, representative of at least 8 individual mice per group. **B** Representative flow cytometric analysis of the splenic B cell compartments Fo.B (CD23^+^/CD21^+^), MZ.B (CD23^-^/CD21^+^) and CD21^lo^.B (CD23^-^/CD21^lo^) after 14 days of treatment with the indicated pellets (left panel). Pie charts in the right panel show average percentages of Fo.B, MZ.B and CD21^lo^.B cells, representative of at least 8 individual mice. **C** Quantification of both percentages (left) and absolute cell numbers (right) of Fo.B cells (blue) and MZ.B cells (red) in murine spleens after 14 days of treatment with Sham (n = 8), Dex (n = 8) and Pred (n = 10) pellets. Mean ± SD. Statistical significance was calculated by applying the ordinary one-way ANOVA or the Kruskal-Wallis test, respectively. **D** Absolute cell numbers of Fo.B cells (blue) and MZ.B cells (red) in murine spleens after 14 days of treatment with Dex (n = 8) and Pred (n = 10) pellets as shown in **C**, normalized to the respective cell numbers in spleens from Sham-treated mice (n = 8). Mean ± SD. Statistical significance was calculated by applying the one sample t test or a mixed effects analysis, respectively. **E** Representative flow cytometric analysis of plasma cells in spleens from mice of the indicated treatment groups (left panel), pre-gated on single viable CD19^+^ cells and quantification of percentages. Sham: n = 8; Dex: n = 7; Pred: n = 9; mean ± SD. Statistical significance was calculated by applying the Kruskal-Wallis test, respectively. **F** Serum IgM & IgG concentrations in mice of the indicated treatment groups determined by ELISA. Sham: n = 7; Dex: n = 8; Pred: n = 9; mean ± SD. Statistical significance was calculated by applying the ordinary one-way ANOVA or the Kruskal-Wallis test, respectively.

FACS staining confirmed the correlation between IgD expression and B cell subpopulations, with follicular B cells showing elevated IgD expression and marginal zone B cells exhibiting reduced IgD expression (Fig. 7B & Fig. S11A). Although total cell numbers decreased across all B cell subpopulations, the reduction in follicular B cells was much more pronounced than that in marginal zone B cells, further highlighting the greater sensitivity of IgD^high^ B cells to GR agonist treatment (Fig. 7C-D).

Since we previously observed that GR agonist treatment enhances activation and promotes plasma cell differentiation *in vitro*, we sought to determine whether the ratio of plasma cells was similarly increased *in vivo* in mice implanted with GC release pellets. Although elevated expression of activation markers CD69 and CD86 was not detectable anymore after 14 days of treatment with GR agonists (Fig. S11B), the ratio of plasma cells, identified by CD138 expression, was elevated in proportion to the potency of the GR agonist treatment (Fig. 7E).

Additionally, IgM serum concentrations were significantly increased in correlation with the strength of the GR agonist effect (Fig. 7F). These findings are consistent with the *in vitro* results, suggesting that IgD-expressing B cells are particularly sensitive to GC-induced cell death, whereas MZ-like B cells, which may have regulatory functions, are relatively resistant. Together with the relatively reduced MZ population, despite the overall increase in total B cells, in the conditional GR knockout (Fig. 1C), our data suggest that GC control the differentiation and survival of B cells in BCR-dependent manner.

## Discussion

Our study highlights the critical role of glucocorticoid receptor (GR) signaling in regulating B cell survival, differentiation, and immune function. Using both *in vivo* and *in vitro* models, we demonstrated that B cells with high IgD expression are particularly sensitive to GC-induced cell death, whereas IgD^low^ (IgM^high^) B cells are relatively resistant, potentially acquiring a regulatory phenotype through increased IL-10 expression. These findings provide important insights into how GCs modulate immune cell populations, thereby supporting and expanding existing literature on immunosuppressive effects elicited by GCs.

GR signaling has long been recognized for its potent immunosuppressive and anti-inflammatory effects, with mechanisms that include apoptosis induction, regulation of cytokine production, and alteration of immune cell differentiation. Surprisingly, we observed that exposure to GCs promotes activation and plasma cell differentiation suggesting that B cells do not merely die by apoptosis but undergo terminal differentiation.

In our study, continuous GR agonist treatment selectively depleted IgD^high^ follicular B cells, which are essential for germinal center responses, while sparing IgM^high^ marginal zone-like B cells. This finding aligns with previous work, which described the ability of the GR to induce apoptosis in various immune cells, including T and B lymphocytes (Gruver-Yates et al., 2014; Wang et al., 2006; Zen et al., 2011). The selective depletion of IgD^high^ B cells may reflect the differential sensitivity of these cells to GR-induced death signals, which we observed both *in vivo* and *in vitro*.

Interestingly, while we did not detect significant differences in the expression of classical pro- or anti-apoptotic genes between IgD^high^ and IgM^high^ B cells, we found that the latter exhibited upregulation of genes such as *Il2rb* and *Cd28* (Table S1, positions 29 and 54, respectively). CD28 signaling has been shown to protect T cells from GC-induced apoptosis, as described by Erlacher et al. (Erlacher et al., 2005), and our data suggest a similar mechanism may play a role in B cells. The upregulation of CD28 in IgM^high^ B cells may enhance their resistance to GC-induced cell death, particularly when activated via TLR signaling pathways, as seen in our CpG and LPS stimulation experiments. This protective effect of CD28 and TLR activation echoes previous studies in T cells, where TCR and CD28 signaling antagonize GC-mediated apoptosis.

Our results further indicate that the activation state of B cells plays a crucial role in determining their sensitivity to GC-induced apoptosis. Resting B cells were highly susceptible to GR-mediated cell death, but TLR activation significantly improved survival. This is consistent with previous findings suggesting that TLR activation can protect plasmacytoid dendritic cells from GC-induced apoptosis (Lepelletier et al., 2010).

In B cells, TLR signaling not only enhances survival but also promotes the differentiation of IL10-producing regulatory B cells (B10 cells), as reported previously (Lepelletier et al., 2010). These regulatory B cells, which we found to be enriched among IgM^high^ populations, are crucial for controlling inflammation and maintaining immune homeostasis through their secretion of anti-inflammatory cytokines like IL-10. The fact that treatment with GR agonists, in the presence of TLR stimulation, selects B cells with high IgM expression and increased IL-10 production, highlights a potential mechanism through which GCs might selectively enrich regulatory cell populations. It is likely that this enrichment is due the sensitivity of IgD^high^ (IgM^low^) B cells to GC-induced cell death. This finding is in agreement with published work, demonstrating that GCs promote the survival of anti-inflammatory macrophages by activating specific signaling pathways (Barczyk et al., 2010). In our study, the selective survival of IgM^high^ B cells with increased IL-10 expression suggests that a similar process may occur in B cells, where GR signaling promotes the enrichment of cells that might contribute to the resolution of inflammation.

Our study also suggests that the GR may exert its effects through both genomic and non-genomic mechanisms. The lack of detectable changes in pro-apoptotic genes like *Bim* or *Puma* in IgD^high^ B cells despite their sensitivity to GR-induced apoptosis suggests that GR may modulate cell death through non-transcriptional pathways. This is supported by previous work showing that the GR can interact with signaling molecules such as Lck and Fyn in T cells, disrupting key signal transduction pathways that are essential for cell survival (Lowenberg et al., 2006). While our transcriptome analysis did not reveal direct involvement of these pathways in B cells, the upregulation of CD28 in IgM^high^ cells hints at a broader role for GR in modulating B cell signaling networks.

Moreover, the role of TLR signaling in protecting B cells from GC-induced apoptosis underscores the importance of innate immune signals in shaping the adaptive immune response during GC exposure. The differential kinetics of CpG and LPS activation, with CpG inducing faster and more robust activation, further supports the idea that the timing and strength of activation signals are critical for determining B cell fate under GC treatment. The delayed GR agonist treatment after LPS activation, which resulted in enhanced survival and plasma cell differentiation, highlights the potential for therapeutic strategies that modulate the timing of immune activation in conjunction with GC therapy.

In conclusion, our findings provide important insights into the selective effects of GR signaling on B cell subpopulations. By promoting the survival of IL-10-producing IgM^high^ B cells and depleting IgD^high^ follicular B cells, GR signaling may contribute to immune regulation and the resolution of inflammation. These results have broad implications for understanding the role of GCs in immune homeostasis and may inform the development of targeted therapies for autoimmune diseases and other inflammatory conditions.

## Author contributions

KA, MA and CS performed mouse analyses, *in vitro* experiments, ELISA and flow cytometric measurements and analyses. KA carried out ELISpot analyses, MY analyzed deep sequencing data. SV, FG and JT performed the *in vivo* treatment with GR agonist-release pellets, provided the GR floxed mice and scientific input. HJ supervised the work and proposed the experiments. KA and CS prepared the figures. HJ and CS designed the study, and wrote the manuscript. All co-authors read and discussed the manuscript.

## Acknowledgements

This work was supported by the DFG through SFB1279 “Exploiting the Human Peptidome for Novel Antimicrobial and Anticancer Agents” to HJ (project B03), SFB1506 “Aging at Interfaces” to HJ (project B05) and JT (project C05 and 450627322). Further project funding by the DFG was granted to SV and FG (project 497680553), SV (project 537908488), JT (project 505870049) and HJ (project 326914434).

## Disclosure of conflicts of interest

The authors declare no competing financial interests.

## Experimental procedures

### Mice

*GR^f/f^* mice (Nr3c1tm2Gsc/J, (Tronche et al., 1999)) were crossed to mb1-cre (CD79a-cre) transgenic mice (Hobeika et al., 2006) in order to achieve B cell-specific cre-mediated deletion in early pro-B cells. C57BL/6J WT mice were bred and housed in the animal facility of Ulm University under specific-pathogen-free conditions. 8 – 30 weeks old male and female mice were used in this study.

GC-releasing pellets (innovative research of america) with Dexamethasone (3.5 mg/pellet) or prednisolone (0.35 mg/pellet) were transplanted into 11 – 22 weeks old male mice. Animal experiments were performed in compliance with license TVA1315 for animal testing at the responsible regional board Tübingen, Germany. All animal experiments were done in compliance with the guidelines of the German law and were approved by the Animal Care and Use Committees of Ulm University and the local government.

### Cell culture

Mature splenic B cells were purified from murine spleens by MACS-based negative selection (B cell isolation kit, mouse - Miltenyi Biotec) according to the instructions provided by the manufacturer. Mature splenic B cells were cultivated in Iscove’s medium (Sigma-Aldrich) containing 10% heat-inactivated FCS (PAN-Biotech), 2 mM L-glutamine (Gibco), 100 U/mL penicillin (Gibco), 100 U/mL streptomycin (Gibco), and 50 μM β-mercaptoethanol (Gibco) at a density of 1 – 2.5 x 10^6^ cells/mL per 6-well (Greiner). Cells were treated either with 2.5 µg/ml LPS (Lipopolysaccharides from Escherichia coli O111:B4, Sigma-Aldrich), 2 µM CpG-ODN #1826 (biomers) or 25 nM Dexamethasone or Prednisolone (Selleckchem), respectively.

### Flow cytometry

Cell suspensions were blocked with α-CD16/CD32 Fc-Block and stained by standard procedures with antibodies enlisted in table 1. Viable cells were distinguished from dead cells by counter-staining with Fixable Viability Dye eFluor 450 or 780 (eBioscience, Thermo Fisher Scientific). For detection of IL-10 secretion, cells were treated with 1 µL/mL GolgiStop (BD Bioscience) for 4 hours before fixation. Intracellular flow cytometry staining was performed by using FIX&PERM Cell Fixation and Permeabilization Kit (ADG Nordic MUbio).

Biotin was detected by using streptavidin Qdot605 (Molecular Probes - invitrogen) Cells were acquired at a FACS Canto II or FACS Lyric flow cytometer (BD Bioscience) and analysis was done by using FlowJo software version 10 (TreeStar). If not indicated otherwise numbers in the dot plots indicate percentages in the respective gates while numbers in histogram plots state the mean fluorescence intensity (MFI).

**Table S1.** Antibodies used in flow cytometric analyses.

| Antigen | Conjugate | Clone | Company |
| --- | --- | --- | --- |
| CD1d | APC | 1B1 | BioLegend |
| CD5 | eFluor 450, PE or PE-Cy-7 | 53-7.3 | eBioscience |
| CD19 | PerCP-Cy5.5 | 1D3 | BD Biosciences |
| CD19 | Biotin | 6D5 | SouthernBiotech |
| CD19 | eFluor 450 | eBio1D3 | eBioscience |
| CD21/CD35 | APC or Biotin | 7E9 | BioLegend |
| CD23 | PE | B3B4 | BD Biosciences |
| CD28 | PE | 37.51 | BioLegend |
| CD43 | Biotin | S7 | BD Biosciences |
| CD45R (B220) | PE-Cy7 or<br>PerCP-eFluor 710 | RA3-642 | eBioscience |
| CD69 | APC or Biotin | H1.2F3 | eBioscience |
| CD86 | PE-Cy7 | GL1 | BioLegend |
| CD93 | APC | AA4.1 | BioLegend |
| CD122 | APC | 5H4 | BioLegend |
| CD138 | PE | DL-101 | eBioscience |
| CD138 | Brilliant Violet 421 | 281-2 | BioLegend |
| GR | Alexa Fluor 647 | D8H2 | Cell Signaling Technology |
| Isotype ctrl | Alexa Fluor 647 | DA1E | Cell Signaling Technology |
| IgD | FITC or Biotin | 11-26 | SouthernBiotech |
| IgD | APC | 11-26 | eBioscience |
| IgM | FITC | polyclonal | SouthernBiotech |
| IgM | eFluor 450 | eB121-15-F9 | eBioscience |
| IgG<sub>1</sub> | Biotin | A85-1 | BD Biosciences |
| IgG<sub>2a</sub> | Biotin | R19-15 | BD Biosciences |
| IgG<sub>2b</sub> | Biotin | R12-3 | BD Biosciences |
| IgG<sub>3</sub> | Biotin | R40-82 | BD Biosciences |
| Ig-κLC | Alexa Fluor 647 | polyclonal | SouthernBiotech |
| Ig-λLC | PE | polyclonal | SouthernBiotech |
| IL-10 | PE | JES5-16E3 | BioLegend |
| IL-10 Isotype<br>ctrl | PE | RTK4530 | BioLegend |

### Enzyme-linked immunosorbent assay

96-well maxisorp plates (NUNC) were coated either with 10 µg/mL polyclonal α- mouse IgM or IgG antibody (SouthernBiotech), respectively, and blocked with PBS (Gibco) containing 1% BSA. Dilutions of mouse IgM or IgG antibodies (Southern Biotech) with given concentrations were used as standards. Murine serum was pre-diluted 1 : 200 in blocking buffer and applied to the plate in duplocates for 1 : 3 serial dilutions. The concentration of IgM or IgG antibodies in the sera was determined by detection with alkaline phosphatase-labeled α-mouse IgM or IgG (Southern Biotech), respectively. P-nitrophenylphosphate (Genaxxon) in diethanolamine-buffer was added and data were acquired at 405 nm using a Multiskan FC ELISA plate reader (Thermo Fisher Scientific).

### ELISpot

Murine splenic cells applied in the ELISpot assay were FACS-purified by using a FACS Aria IIu cell sorter (BD), as shown previously (Setz et al., 2018). For detection of IgM 10 x 10^3^, for IgG 50 - 100 x 10^6^ cells were plated in triplicates. The assay was performed for IgM and IgG (ELISpot Basic ALP, Mabtech) according to manufacturer’s instructions and pictures were acquired and analyzed by using an Astor ELISpot reader (Mabtech).

### RNA sequencing

Splenic B cells were purified and cultivated as described in the cell culture section. After three days of treatment with LPS, B cells were FACS-stained with anti-IgM and IgD antibodies and sorted as shown in **Fig. S9** depending on their IgD expression into IgD^high^ and IgD^low^ B cell fractions by using a FACS Aria IIu cell sorter (BD). Transcriptome analysis was performed by GENEWIZ Germany GmbH.

RNA sequencing (RNA-seq) data were processed and analyzed using a series of steps starting with quality control and preprocessing. Raw paired-end fastq files were trimmed for adapter sequences and low-quality reads using Trimmomatic (version 0.39), with a sliding window approach and a minimum read length of 75 nucleotides. The cleaned reads were aligned to the Mus musculus GRCm39 reference genome using Hisat2 (version 2.2.1), and the resulting alignments were converted to sorted BAM files using Samtools (version 1.9). Gene-level counts were generated using FeatureCounts from Subread (version 2.0.6).

For differential gene expression analysis, the count data were imported into edgeR (version 3.32.1). To minimize the influence of lowly expressed genes, a filtering step was performed using the counts per million (CPM) values. Genes with CPM values greater than 8 in at least 6 samples were retained for further analysis. This threshold effectively removed genes with insufficient read counts across samples. Following this, the data were normalized using TMM (Trimmed Mean of M-values) normalization to account for differences in library size.

Differential expression was assessed by fitting a generalized linear model (GLM) and performing likelihood ratio tests (LRT) on group comparisons. Genes with an absolute log-fold change (logFC) ≥ 1.5 (corresponding to a fold change of ≥ 2) and an FDR < 0.05 were considered differentially expressed. To further refine the analysis, the glmTreat function was used to enforce the fold-change threshold of 1.5

### Statistical analysis

Graphs were created and statistical analysis was performed by using GraphPad Prism (Version 10) software. The numbers of individual replicates or mice (n) is stated in the figure legends as well as the tests applied to calculate statistical significance among observed differences. P values < 0.05 were considered to be statistically significant (n. s. = not significant; * p ≤ 0.05; ** p ≤ 0.01; *** p ≤ 0.001; **** p ≤ 0.0001).

## Notes

### Competing Interest Statement

The authors have declared no competing interest.

