## Supplementary Figures S1 - 12 for "IgD-Expressing Mature B Cells Exhibit Enhanced Sensitivity to Glucocorticoid-Induced Cell Death"

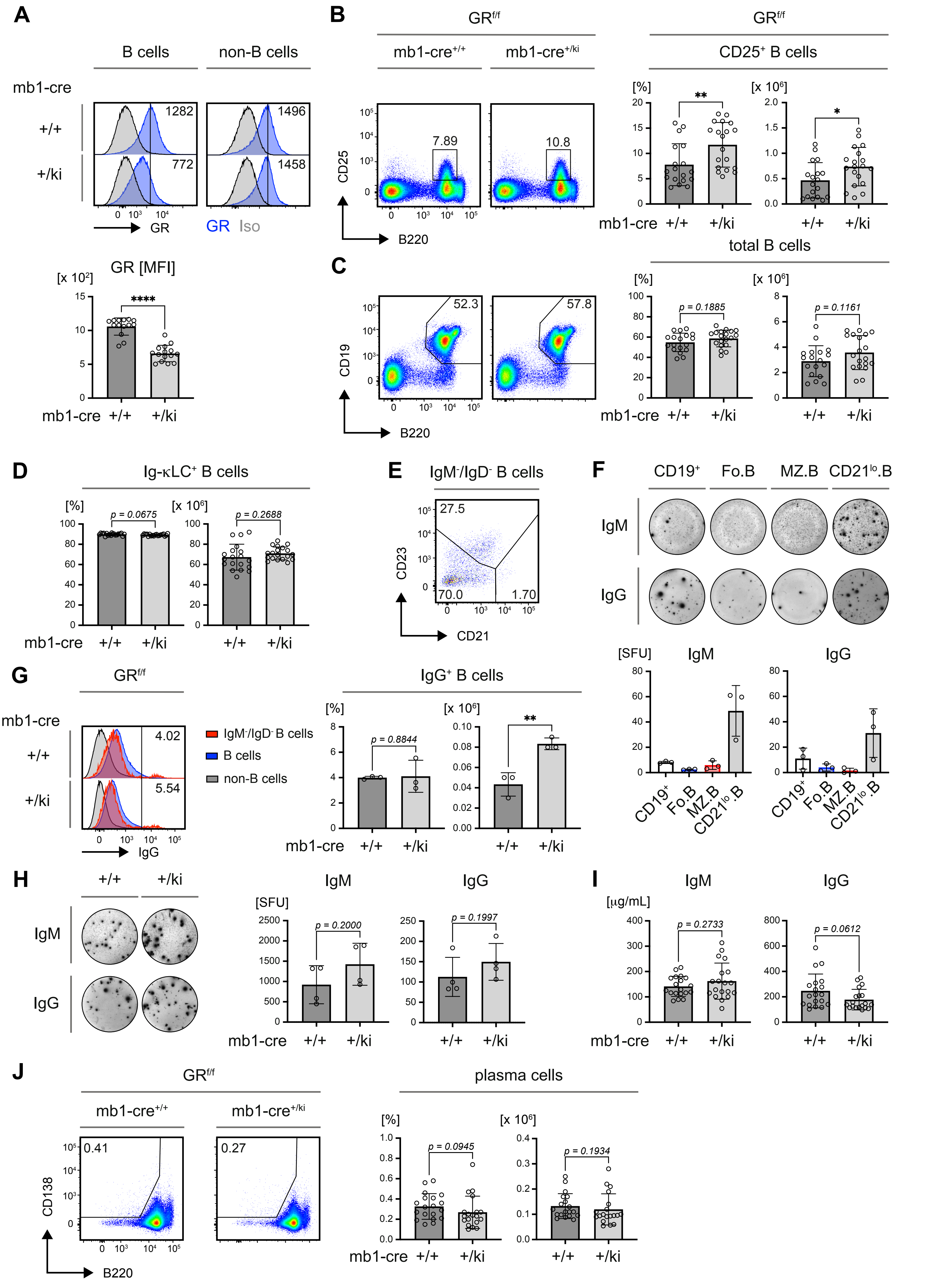

**Figure S1 | B cell-specific GR deletion alters splenic B cell subpopulations**

**Figure S1 | B cell-specific GR deletion alters splenic B cell subpopulations (related to main Figure 1)**

Phenotype analyses of B cell populations in 8 weeks old glucocorticoid receptor (GR) *GR$^{f/f}$ x mb1-cre* mice of the indicated genotype, n = 19 in each group, if not indicated otherwise; mean ± SD. Flow cytometric data were pre-gated on B cells as shown in **Fig. 1A** (left panel). Statistical significance was calculated by applying either the unpaired t test or Mann-Whitney-U test depending on the distribution of data points.

**A |** Representative comparison of glucocorticoid receptor (GR) expression between splenic B and non-B cells from mice of the indicated genotypes (top panel, values in histogram plots indicate the respective mean fluorescence intensities (MFI)) and MFI quantification (bar diagrams, bottom). *GR$^{f/f}$ x mb1-cre$^{+/+}$*: n = 14; *GR$^{f/f}$ x mb1-cre$^{+/ki}$*: n = 15; mean ± SD. Statistical significance was calculated by applying the Mann-Whitney-U test.

**B |** Representative flow cytometric analysis (left panel) and quantification of both percentages (bar diagrams, left) and absolute cell numbers (bar diagrams, right) of small pre-B cells identified by expression of CD25 in the bone marrow from mice of the indicated genotypes. *GR$^{f/f}$ x mb1-cre$^{+/+}$*: n = 18; *GR$^{f/f}$ x mb1-cre$^{+/ki}$*: n = 19; mean ± SD. Statistical significance was calculated by applying the Mann-Whitney-U test or the unpaired t test, respectively.

**C |** Representative flow cytometric analysis (left panel) and quantification of percentages (bar diagrams, left) and absolute cell numbers (bar diagrams, right) of total B cells identified by expression of CD19 and B220 in the bone marrow from mice of the indicated genotypes. *GR$^{f/f}$ x mb1-cre$^{+/+}$*: n = 18; *GR$^{f/f}$ x mb1-cre$^{+/ki}$*: n = 19; mean ± SD. Statistical significance was calculated by applying the unpaired t test.

**D |** Quantified percentages of intracellular Igκ-LC$^{+}$ B cells (left panel) and absolute cell numbers (bar diagrams, panel) in the spleens of mice from the indicated genotypes.

**E |** Representative flow cytometric analysis of CD21 and CD23 expression in IgM$^{-}$/IgD$^{-}$ B cells derived from a *GR$^{f/f}$ x mb1-cre$^{+/ki}$* mouse, pre-gated as shown in **Fig 1F**.

**F |** Representative ELISpot data of IgM & IgG secreting cells in spleens of indicated B cell populations (left panel) purified from wild-type (WT) mice and quantification of spot forming units (SFU, right panel). n = 3; mean ± SD.

**G |** Representative flow cytometric analysis of IgG$^{+}$ B cells within the population of IgM$^{-}$/IgD$^{-}$ splenic B cells (left panel) and quantification of percentages (bar diagrams, left) and absolute cell numbers (bar diagrams, right). n = 3; mean ± SD. Statistical significance was calculated by applying the unpaired t test.

**H |** Representative ELISpot data of IgM & IgG secreting cells in FACS-purified splenic IgM$^{-}$/IgD$^{-}$ B cell populations derived from *GR$^{f/f}$ x mb1-cre* mice of the indicated genotypes (left) and quantification of SFU (right), n = 4; mean ± SD.

**I |** Serum IgM (top) and IgG (bottom) concentrations in mice of the individual genotypes determined by ELISA.

**J |** Representative flow cytometric analysis of plasma cells in the spleen (top panel), pre-gated on viable single CD19$^{+}$ cells and quantification of percentages (bar diagrams, bottom left) and absolute cell numbers (bar diagrams, bottom right).

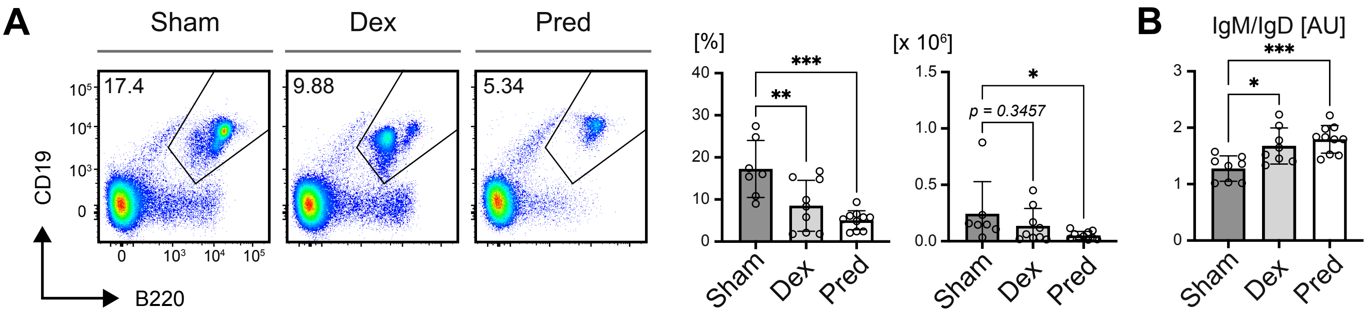


**Figure S2 | Continuous GC-treatment eradicates B cells *in vivo***
**(related to main Figure 2)**

Phenotype analyses of mice transplanted with constant glucocorticoid (GC)-release pellets after 14 days of GC treatment.

**A |** Representative flow cytometric analysis (left panel) and quantification of both percentages (bar diagrams, left) and absolute cell numbers (bar diagrams, right) of total B cells in the bone marrow from mice following 14 days of exposure to Dexamethasone (Dex, n = 9), Prednisolone (Pred, n = 10) or control (Sham, n = 7) pellets. Mean ± SD. Statistical significance was calculated by applying the ordinary one-way ANOVA or the Kruskal-Wallis test, respectively.

**B |** Quantified ratios of IgM/IgD surface expression (displayed as arbitrary units, AU) in splenic B cells from Sham- (n = 8), Dex- (n = 7) and Pred-treated (n = 9) mice. Values used for calculation are displayed in **Fig 2F**. Mean ± SD. Statistical significance was calculated by applying the ordinary one-way ANOVA.

### Figure S2 | Continuous glucocorticoid treatment eradicates B cells *in vivo*

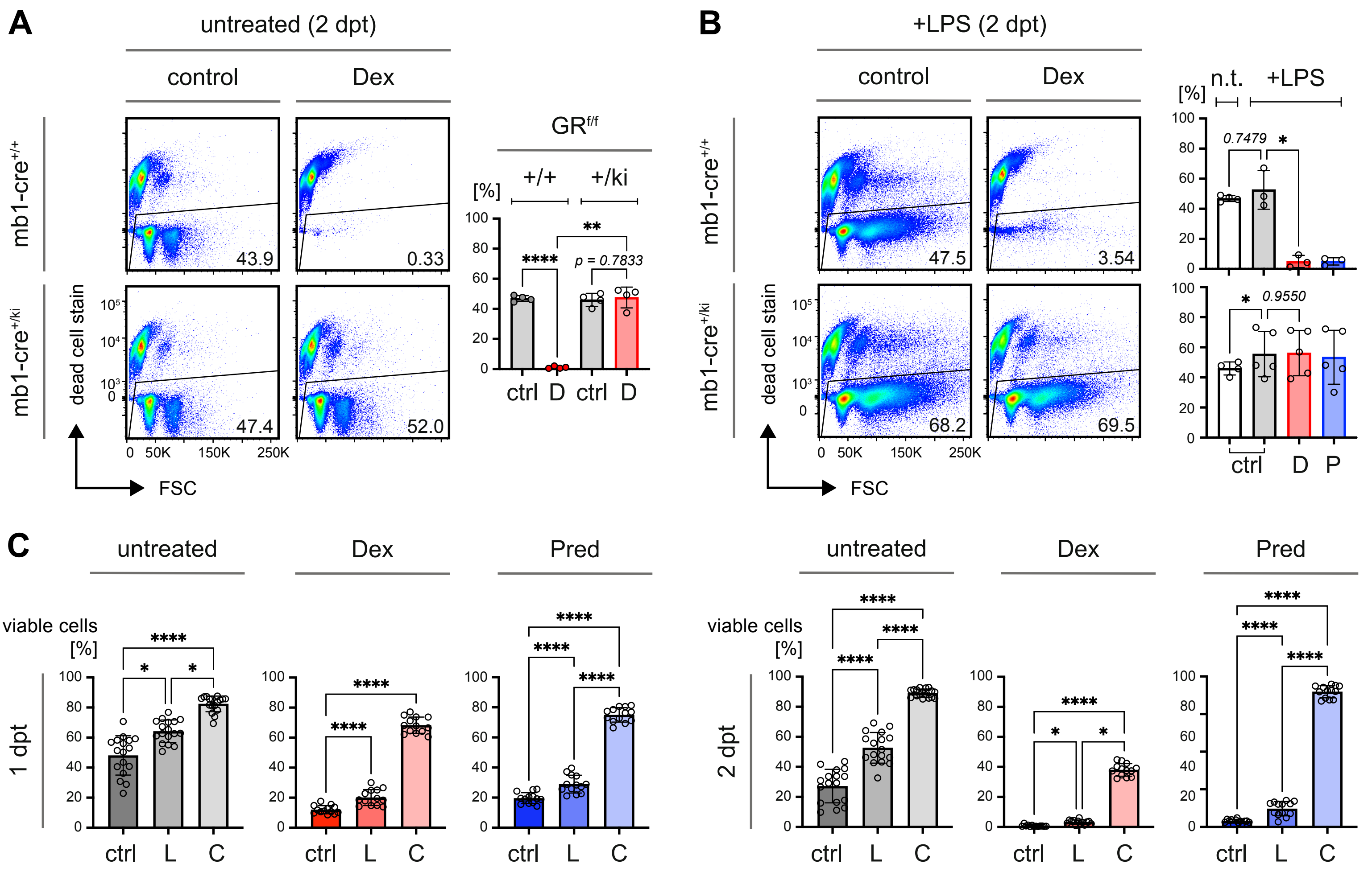


**Figure S3 | Activated mature B cells are more resistant to GR agonist treatment *in vitro* (related to main Figure 3)**

**A - B |** Representative comparison (left) and quantification (right) of viability in splenic B cells from *$GR^{f/f}$ x mb1-cre* mice of the indicated genotypes presence (**B**, n = 5 for each group and genotype) or absence (**A**, n = 4 for each group and genotype) of 2.5 µg/mL lipopolysaccharide (LPS) upon treatment with 25 nM GR agonists Dex (D) or Pred (P) at 2 days post treatment (dpt) *in vitro* as described in **Fig. 3A**; mean ± SD. Statistical significance was calculated by applying the repeated measures (RM) one-way ANOVA.

**C |** Quantified percentages of viable cells from **Fig. 3B&D** were compared at 1 (left panel) and 2 dpt (right panel) when treatment with GR agonists was performed in presence of either LPS (L, **Fig. 3C**) or CpG (C, **Fig. 3D**) or without pre-stimulation (ctrl, **Fig. 3B**). Control (black, n = 17), Dex (red, n = 13), Pred (blue, n = 17), mean ± SD. Data from mice included in the kinetics shown in **Fig. 3B&D** in which conditions were missing were excluded in this analysis in order to perform a paired statistical test. Statistical significance was calculated by the repeated measures (RM) one-way ANOVA or Friedman test, respectively.

#### Figure S3 | Activated B cells display relative resistance to treatment with GR agonists

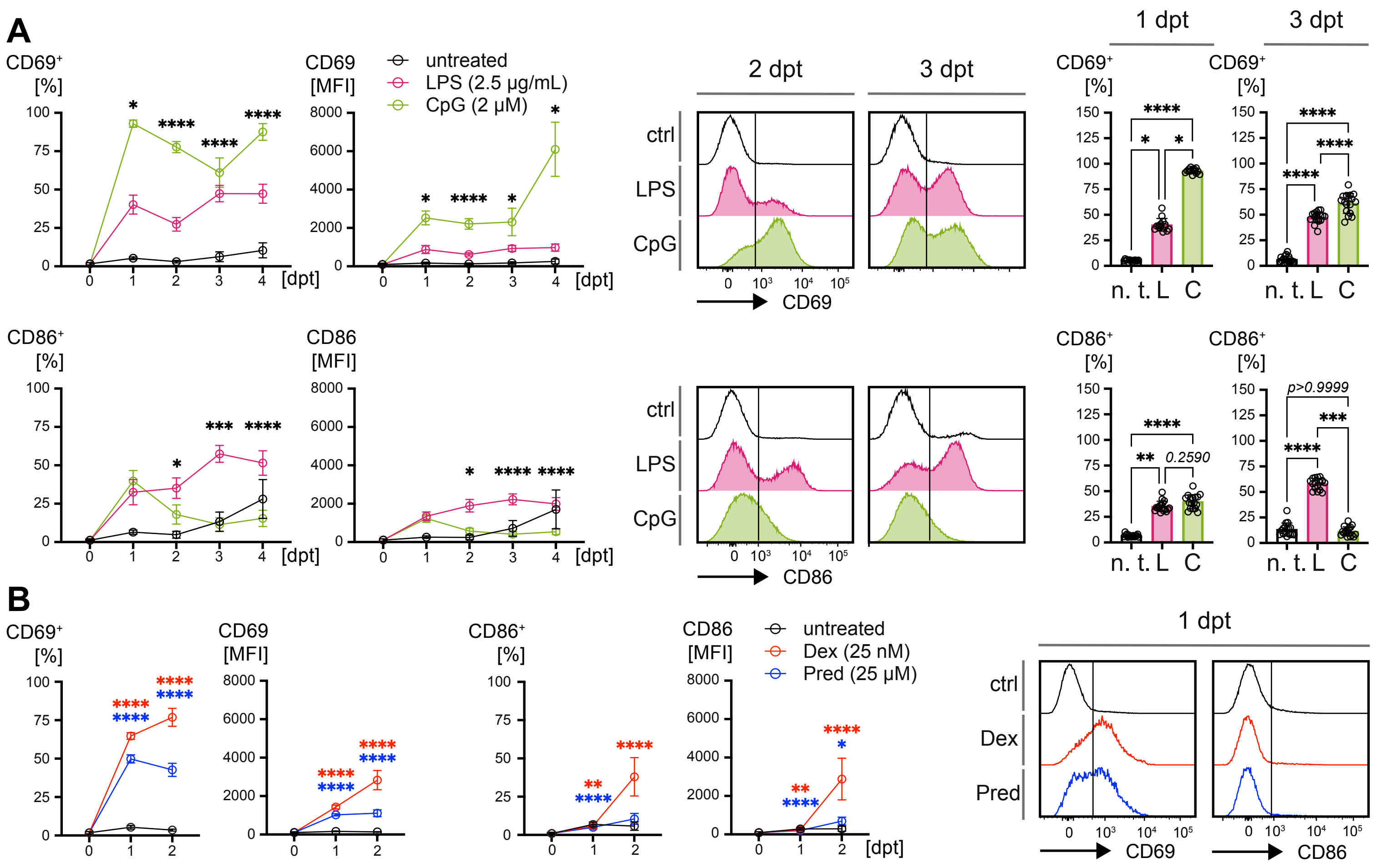


**Figure S4 | GR agonists enhance B cell activation and accelerate terminal differentiation (related to main Figure 4)**

**A |** Kinetics of activation markers CD69 (top panel) and CD86 (bottom panel), determined by flow cytometry in WT splenic B cells following treatment with LPS and CpG, respectively. Representative flow cytometric analyses and bar diagrams on the right-hand side compare percentages of CD69 and CD86 positive cells upon LPS- or CpG-treatment at the indicated time points. n = 17 for all groups and time points, except for 0 dpt (n = 11), mean ± SD. Statistical significance was calculated by applying either the RM one-way ANOVA or the Friedman test, respectively. Asterisks in the kinetics indicate significant differences between cells stimulated either with LPS or CpG.

**B |** Kinetics of activation markers CD69 (left panels) and CD86 (middle panels), determined by flow cytometry in WT splenic B cells following treatment with the GR agonists Dex or Pred (both at a concentration of 25 nM) in absence of LPS or CpG. Histograms on the right-hand side show representative flow cytometric analyses of CD69 and CD86 at 1 dpt. For CD69 day 0: n = 9; day 1: n = 8; day 2: n =11. For CD86 day 0: n = 9; day 1 and 2: n = 11; mean ± SD. Statistical significance was calculated by applying either the RM one-way ANOVA or the Friedman test, respectively. Asterisks in the kinetics indicate significant differences between cells stimulated either with Dex or Pred and untreated controls.

### Figure S4 | GCs enhance B cell activation and accelerate terminal differentiation

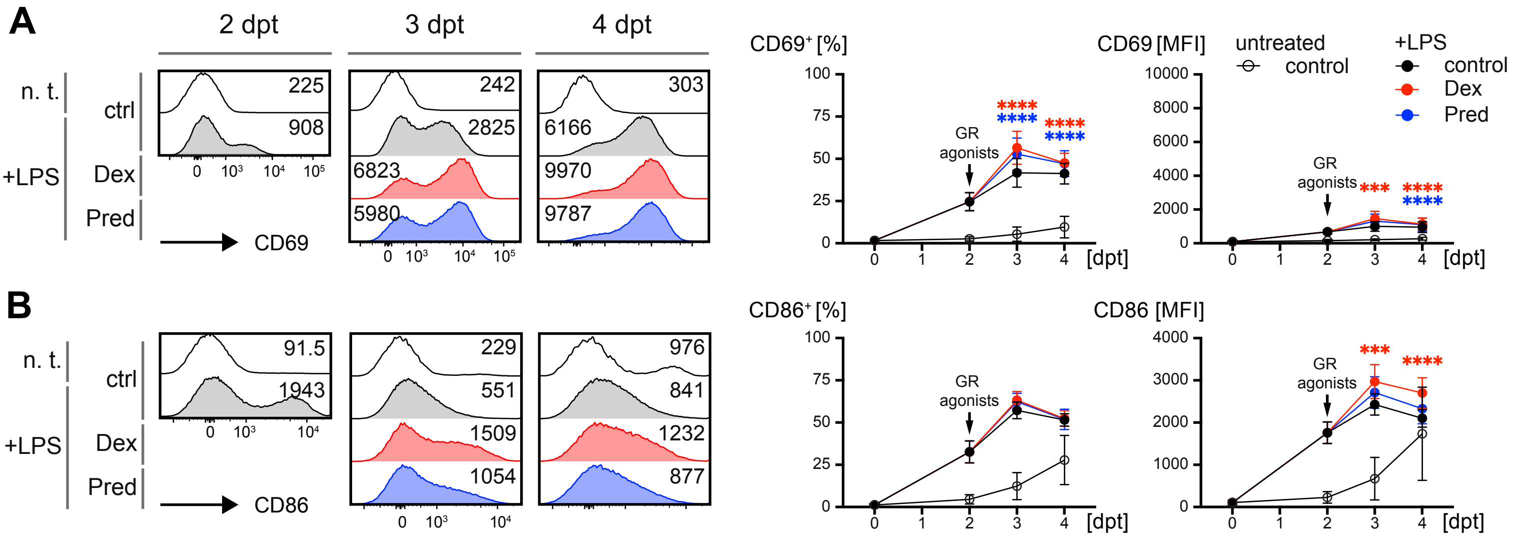


**Figure S5 | CD69$^{hi}$ mature B cells are more robust to treatment with GR agonists (related to main Figure 5)**

**A - B |** Representative flow cytometric analysis of activation markers CD69 (**A**) and CD86 (**B**) in WT splenic B cells at the indicated time points following treatment with GR agonists as described in **Fig. 5A** in presence of LPS. Kinetics beside the histograms compare percentages of positive cells (left) and MFI (right) of CD69 (A) and CD86 (B). Day 0: n = 11; day 2 & 4: n =15; day 3: n =14; mean ± SD. Statistical significance was calculated by applying the paired t test for day 2 and either the RM one-way ANOVA or the Friedman test for assessing differences at days 3 and 4. Asterisks in the kinetics indicate significant differences between cells stimulated either with LPS and cells additionally treated with GR agonists Dex (D, red asterisks) or Pred (P, blue asterisks).

**Figure S5 | Delayed GR agonist treatment after LPS activation improves B cell survival**

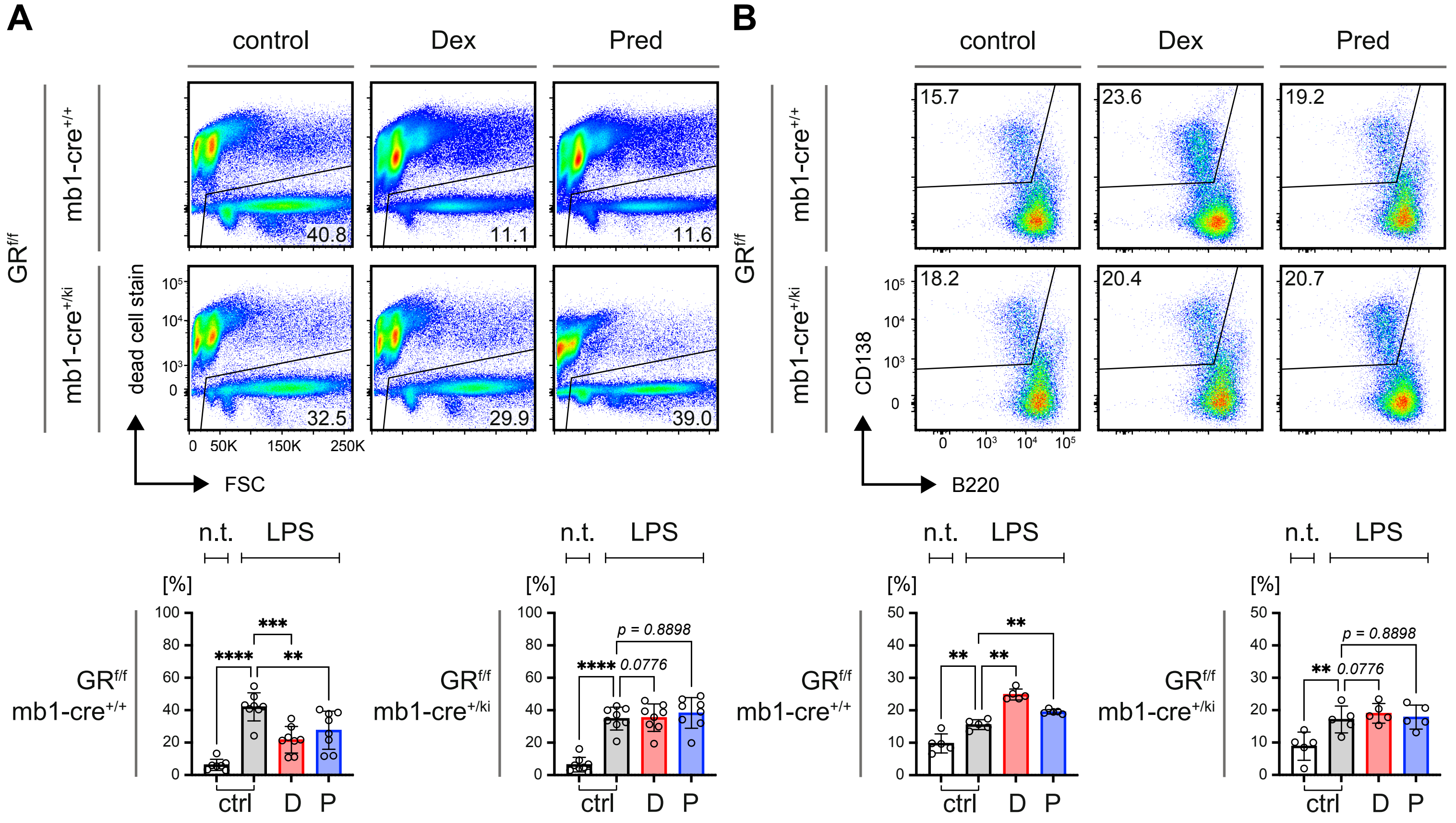


**Figure S6 | GR-deficient B cells are more resistant to treatment with GR agonists (related to main Figure 5)**

**A |** Representative flow cytometric analyses of cellular viability upon GC-treatment of purified splenic B cells derived from *GR$^{f/f}$ x mb1-cre* mice of the indicated genotypes at 3 dpt *in vitro* as described in **Fig. 5A**. Bar diagrams below show quantified of percentages of viable cells, respectively. Mean ± SD. Statistical significance was calculated for the *GR$^{f/f}$ x mb1-cre$^{+/+}$* dataset (left, n = 7 for n. t. ctrl, n = 8 for all other groups) by applying the mixed-effects analysis with Šídák's multiple comparisons test and for the *GR$^{f/f}$ x mb1-cre$^{+/ki}$* dataset (left, n = 9 for each group) by applying the RM one-way ANOVA, respectively.

**B |** Representative flow cytometric analyses of plasma cell differentiation in vitro upon GC-treatment of purified splenic B cells derived from *GR$^{f/f}$ x mb1-cre* mice of the indicated genotypes at 3 dpt as described in **Fig. 5A**. Bar diagrams below show quantified of percentages of CD19$^{+}$/CD138$^{+}$ cells, respectively. n = 6 for all groups and genotypes, mean ± SD. Statistical significance was calculated by applying the RM one-way ANOVA, respectively.

### Figure S6 | GR-deficient B cells are resistant to GR-agonist treatment

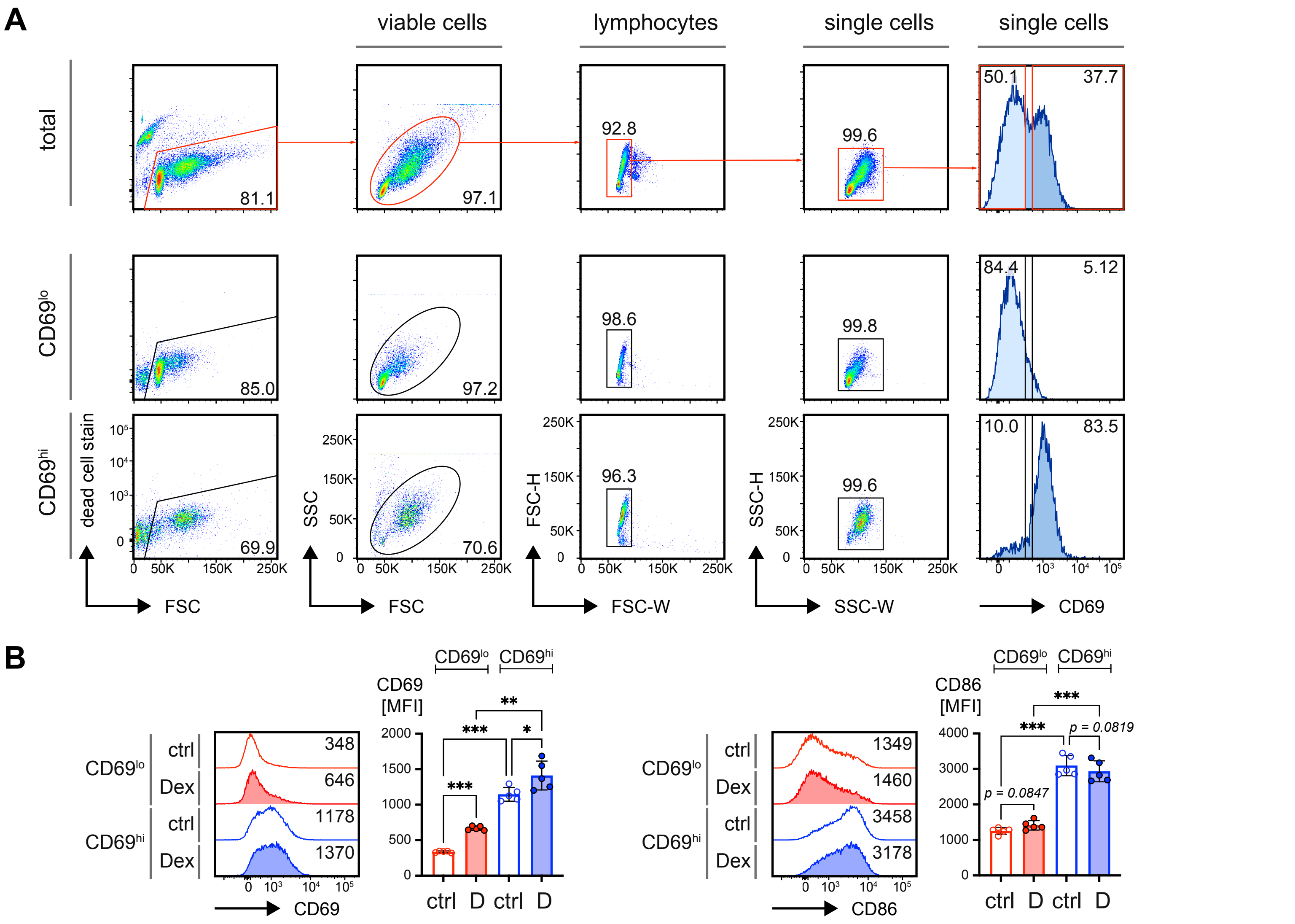


**Figure S7 | Sorting of CD69$^{hi}$ and CD69$^{lo}$ B cells for treatment with GR agonists (related to main Figure 5)**

**A |** FACS-purification of CD69$^{+}$ and CD69$^{-}$ mature splenic B cells from WT mice. Mature splenic B cells were isolated by negative selection via magnetic activated cell sorting (MACS) and treated for 2 days with 2.5 µg/mL LPS. At 2 dpt cells were stained for CD69 and FACS-purified according to the gating strategy, shown in the top row. Subsequently the CD69$^{lo}$ and CD69$^{hi}$ populations were re-analyzed to assess purity. Representative data are shown in the two lower rows.

**B |** Representative flow cytometric analysis of activation markers CD69 and CD86 (left) and quantified MFI (right) in FACS-purified CD69$^{lo}$ and CD69$^{hi}$ mature splenic B cells from WT mice treated overnight in presence (D) or absence (ctrl) of 25 nM Dex, n = 5, mean ± SD. Statistical significance was calculated by applying either the RM one-way ANOVA.

**Figure S7 | Sorting of CD69$^{hi}$ and CD69$^{lo}$ B cells for treatment with GR agonists**

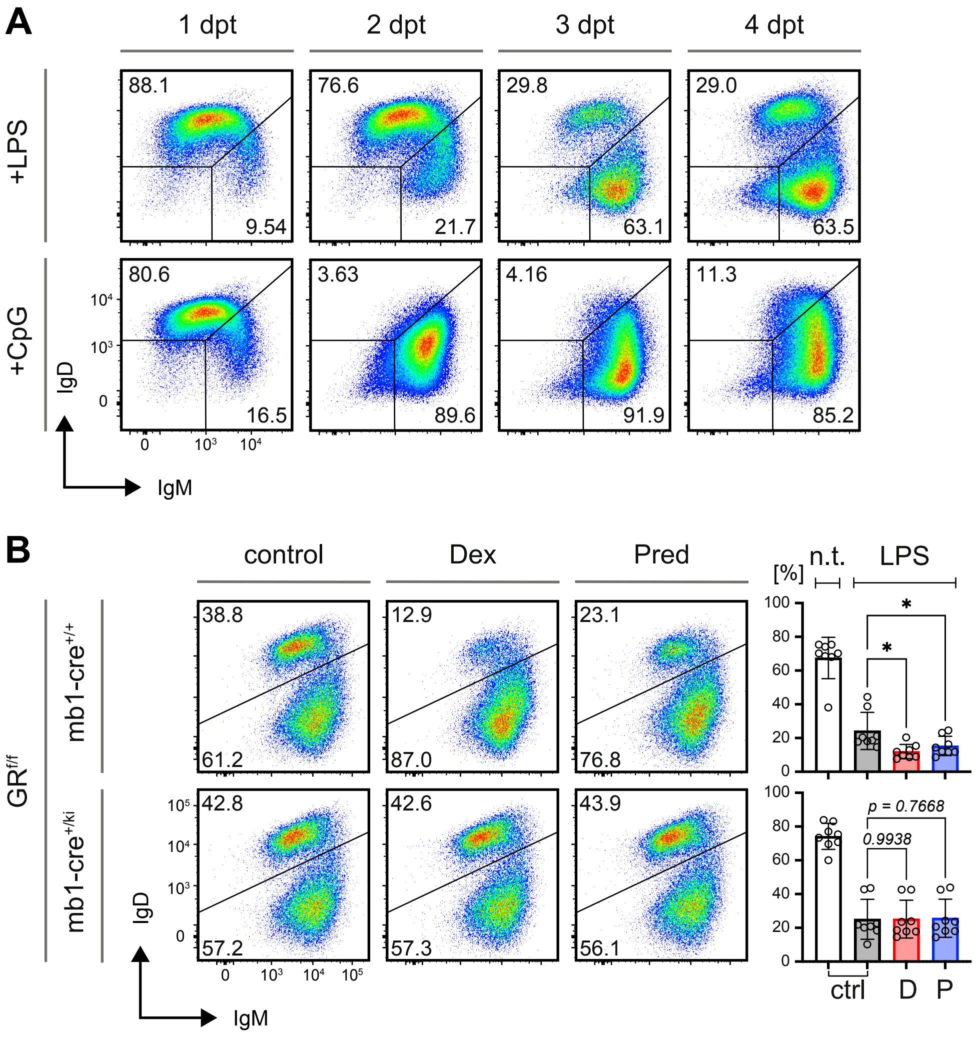


**Figure S8 | IgD BCR expression affects resistance to cell death (related to main Figure 6)**

**A |** Representative flow cytometric analyses of IgM and IgD BCR surface expression, determined at the indicated time points by flow cytometry in WT splenic B cells following treatment with LPS and CpG. Shown data are representative of at least 6 individual mice for all groups and time points, mean ± SD.

**B |** Representative flow cytometric analyses of IgM and IgD surface expression in purified splenic B cells derived from $GR^{f/f}$ *x mb1-cre* mice of the indicated genotypes at day 3 upon stimulation with LPS and treatment with delayed GCs as described in **Fig. 5E**. Bar diagrams show quantified percentages of $IgM^{lo}/IgD^{hi}$ B cells cultivated in presence of LPS and following GC treatment n = 6 for all groups and genotypes, except for $GR^{f/f}$ *x mb1-cre*$^{+/ki}$ n. t./ctrl: n = 5. Statistical significance was calculated by applying the RM one-way ANOVA, respectively.

### Figure S8 | IgD BCR expression affects resistance to cell death

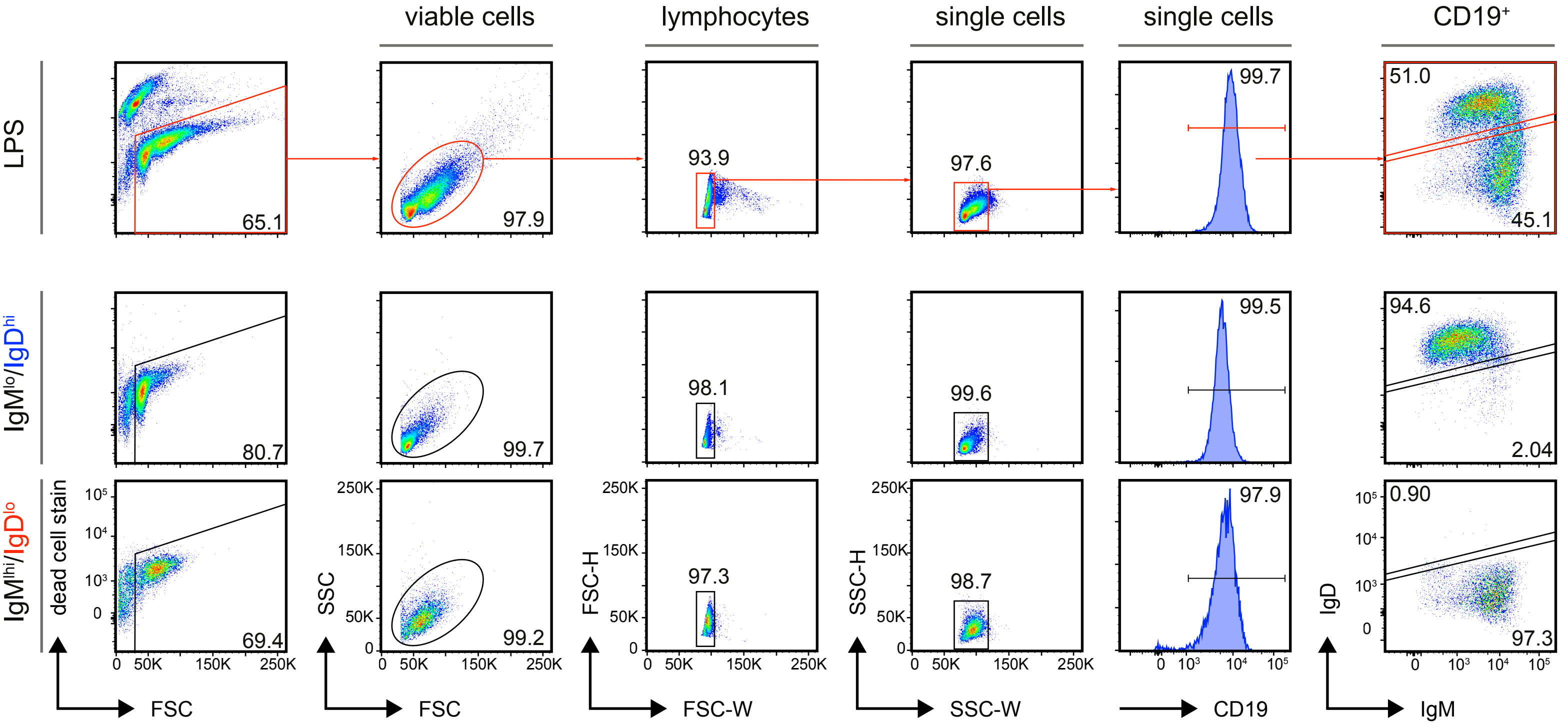


**Figure S9 | Sorting strategies, gating and purity of IgD$^{hi}$ and IgD$^{lo}$ B cells (related to main Figure 6)**

Gating strategy (top row) for FACS-purification of IgM$^{lo}$/IgD$^{hi}$ and IgM$^{hi}$/IgD$^{lo}$ B cell populations and representative purity (bottom rows) of sorted cell populations. B cells from WT mice were purified by negative selection via MACS stimulated with 2.5 µg/mL LPS or left untreated. After 2 days of incubation, cells were FACS-purified to separate IgM$^{lo}$/IgD$^{hi}$ and IgM$^{hi}$/IgD$^{lo}$ populations.

**Figure S9 | Sorting strategies, gating and purity of IgD$^{hi}$ and IgD$^{lo}$ B cells**

A

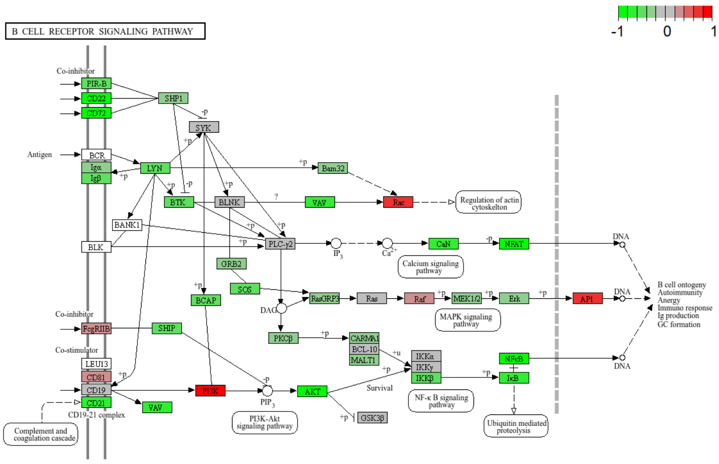


B

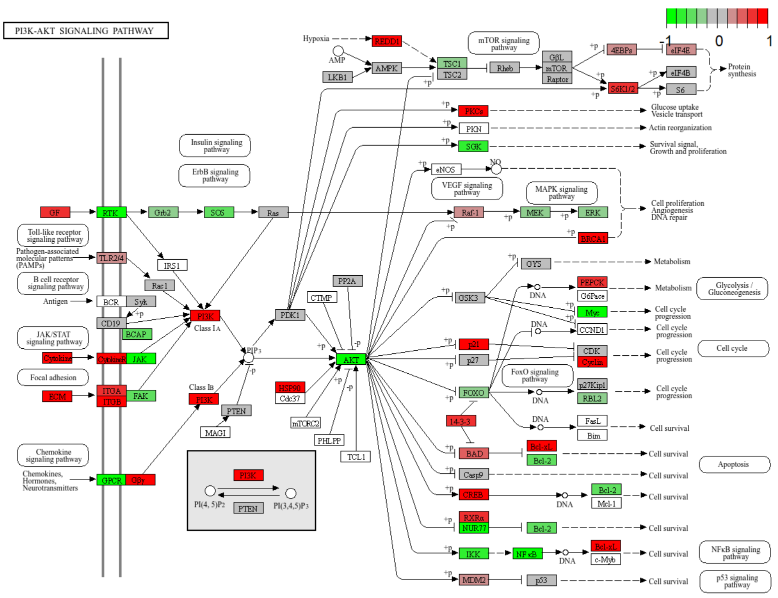


C

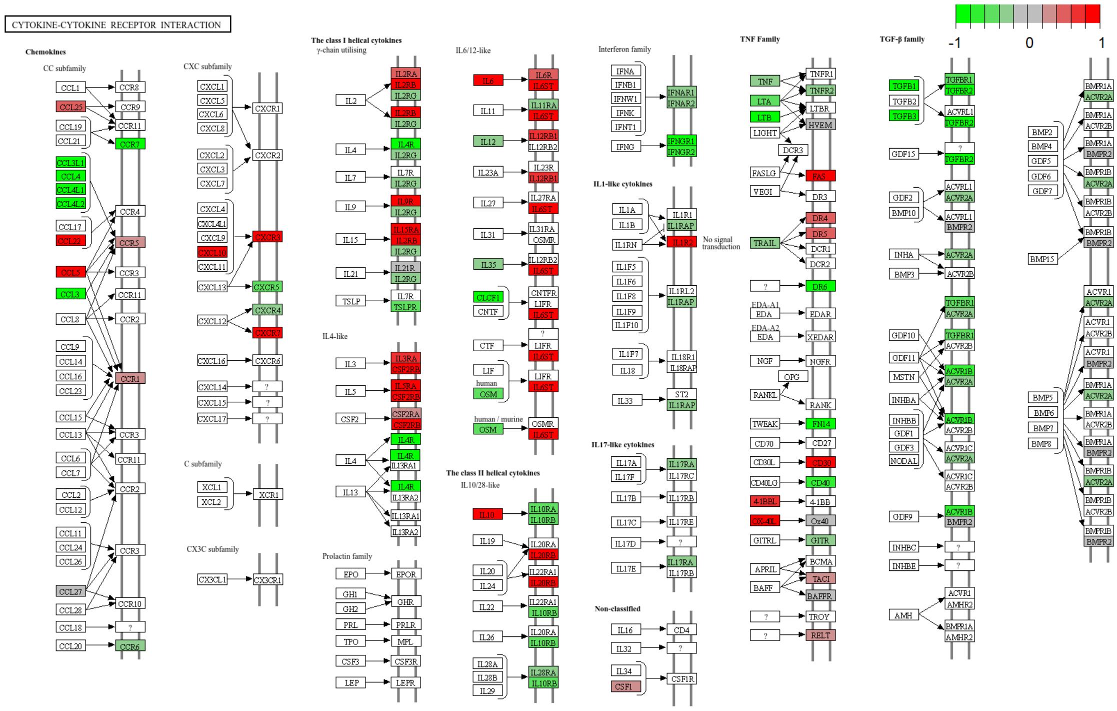


**Figure S10 | KEGG pathway analysis in IgM$^{lo}$/IgD$^{hi}$ and IgM$^{hi}$/IgD$^{lo}$ B cell populations (related to main Figure 6)**

**A |** B cell receptor (BCR) signaling pathway

**B |** Phosphoinositide 3-kinase (PI3K)-protein kinase B (PKB/AKT) signaling pathway

**C |** Cytokine-cytokine recptor interactions

### Figure S10 | KEGG pathway analysis

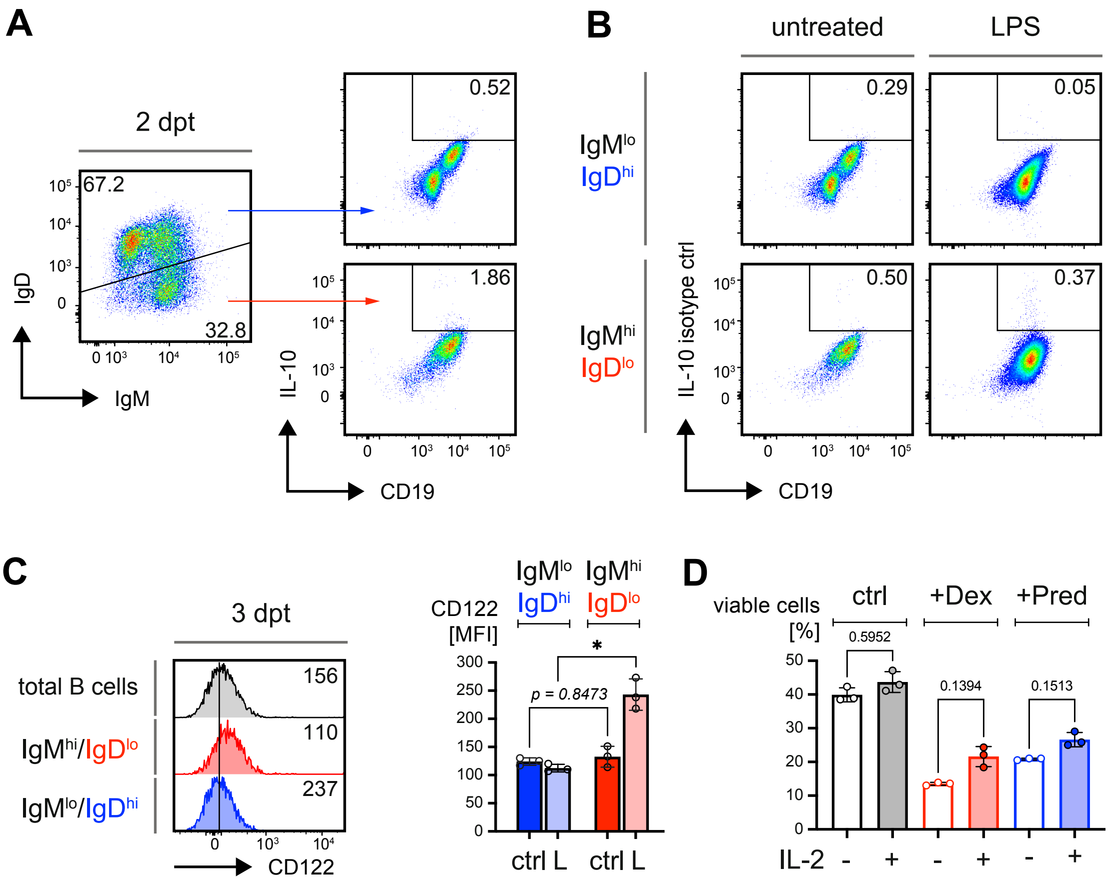


**Figure S11 | IgD BCR expression affects resistance to cell death (related to main Figure 6)**

**A |** Representative flow cytometric analysis of IL-10 expression in untreated IgM$^{lo}$/IgD$^{hi}$ and IgM$^{hi}$/IgD$^{lo}$ splenic B cells at 4 dpt. Mature splenic B cells from WT mice were purified by negative selection via MACS and left untreated.

**B |** Representative flow cytometric analysis of the isotype control used to assess the background of the antibody used for detection of IL-10 in untreated IgM$^{lo}$/IgD$^{hi}$ and IgM$^{hi}$/IgD$^{lo}$ splenic B cells at 4 dpt. Mature splenic B cells from WT mice were purified by negative selection via MACS and left untreated.

**C |** Representative flow cytometric analysis of CD122 expression in purified mature B cells and IgM$^{lo}$/IgD$^{hi}$ and IgM$^{hi}$/IgD$^{lo}$ subpopulations upon treatment with 2.5 µg/mL LPS for 3 days (left). Quantification of CD122 MFI in IgM$^{lo}$/IgD$^{hi}$ and IgM$^{hi}$/IgD$^{lo}$ B cells. n = 3, mean ± SD. Statistical significance was calculated by applying the RM one-way ANOVA.

**D |** Quantified survival of mature B cells in presence and absence of IL-2 during exposure to GR agonists. Mature splenic B cells from WT mice were purified by negative selection via MACS and treated with 2.5 µg/mL LPS. At 2 dpt cells were treated with Dex, or Pred in presence or absence of 0.3 µg/mL IL-2. Percentages of viable cells were quantified by flow cytometry at 4 dpt (2 days following addition of IL-2 and GR agonists). n = 3, mean ± SD. Statistical significance was calculated by applying the RM one-way ANOVA.

### Figure S11 | IgD$^{low}$ B cells show enhanced survival and increased IL-10 expression

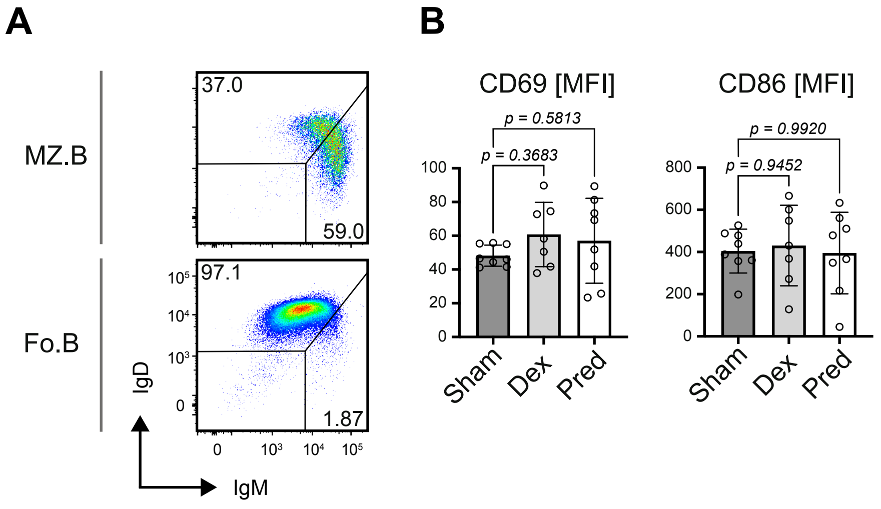


**Figure S12 | Selective effects of GR agonist treatment on IgD expressing B cells *in vivo***
**(related to main Figure 7)**
Phenotype analyses of mice transplanted with constant glucocorticoid (GC)-release pellets after 14 days of GC treatment.
**A |** Representative flow cytometric analysis of IgM and IgD surface expression by follicular (Fo.B) and marginal zone B (MZ.B) cells purified an untreated WT mouse.
**B |** Quantified MFI of CD69 (left) and CD86 MFI in splenic B cells from Sham- (n = 8), Dex- (n = 7) and Pred-treated (n = 8) mice. Mean ± SD. Statistical significance was calculated by applying the ordinary one-way ANOVA.
